# Pinning transition in biofilm structure driven by active layer dynamics

**DOI:** 10.1101/2022.03.21.485164

**Authors:** Ellen Young, Gavin Melaugh, Rosalind J Allen

## Abstract

Surface-attached communities of microbes, known as biofilms, are diverse in their morphologies. Characterising distinct types of biofilm spatial structure, and understanding how they emerge, can shed light on the fundamental biological and biophysical mechanisms involved, and can improve our understanding of evolution in biofilms. Here, we perform long-time individual-based simulations of growing biofilms. We observe distinct types of biofilm spatial structure depending on the parameters, and we classify these into three ‘phases’ according to the behaviour of the active layer of growing cells at the biofilm interface. In the unpinned phase, the biofilm is smooth and the active layer is unbroken with no gaps. In the transiently pinned phase, short-lived gaps in the active layer arise, which can cause local parts of the biofilm interface to pin, or become stationary relative to the moving front. In the pinned phase these ‘pinning sites’ persist, leading to fingering of the biofilm interface. We show that pinning arises due to the dynamical behaviour of active layer gaps, and observe that the relative magnitudes of the active layer thickness and the active layer fluctuations are important in this process. We demonstrate a direct connection between biofilm pinning and interface roughness, and we show that the pinning phase transition is well described by a control parameter that combines the average and standard deviation of the active layer thickness. Taken together, our work suggests a role for active layer dynamics in controlling pinning of the biofilm interface and hence biofilm morphology.

## Introduction

Biofilms, or surface-attached populations of microbes, are diverse in their morphologies. Biofilms grown under flow can be smooth or rough, or even ‘mushroom-shaped’^1–3^, while biofilms on liquid interfaces, and bacterial colonies grown on agar plates, can show intricate wrinkly patterns^4^. In the biophysics community, colonies growing on agar plates have attracted significant interest, with morphologies ranging from circular to fractal-like, and even chiral patterns^5–8^. Characterising distinct types of biofilm or colony spatial structure, and the mechanisms by which they emerge, can lead to better understanding of the biophysics of multicellular growth, as well as self-assembly and non-equilibrium growth processes more generally. Understanding biofilm spatial structure is also a prerequisite for understanding a diverse range of phenomena, including genetic mixing and hence potential for cooperation, the extent of pathogen adhesion, as well as antibiotic penetration and the chances of fixation of antibiotic resistant mutants^9–13^.

There are a number of important debates about what produces different biofilm spatial structures, which is often characterised in terms of the interface roughness, *i*.*e*. the standard deviation of the height. From a mechanistic point of view, it is well established that the interplay between local growth and the nutrient concentration field is important in controlling the roughness of biofilm interfaces^14–16^. Specifically, Dockery and Klapper^15^ showed the existence of a fingering instability in which a local ‘bump’ on the growing interface tends to grow larger, since bacteria in the ‘bump’ have better access to nutrients (diffusing from above) than those in adjacent areas; the growing bump then depletes nutrients from adjacent regions of the interface, further enhancing its growth. More recently, attention has also been focused on the role of mechanical interactions, which arise from cells pushing against each other and the substrate as they grow^17–25^ and may drive a transition from a smooth growing interface to a rough one^17^. Other factors that are likely to be important are the production of exopolymeric substances^19^, immigration of new cells to the biofilm^2,26^ and interaction with local fluid flow fields^27^. Statistical physicists have also attempted to classify different regimes of interface growth in bacterial communities based on the scaling behaviour of their interface roughness, allowing comparison to universal growth laws well known in statistical physics.^13,28–31^.

A key concept in our understanding of biofilm (or colony) growth is that of the active layer (Figure 1), where growth occurs only in a layer of finite thickness at the edge of the biofilm that has access to nutrients, while cells deeper within the biofilm are not able to grow^8,32–34^. This phenomenon is observed in simulations^33,35^, experimental flow cells and colonies^34,36^(colony ref) as well as in *in vivo* samples^34^, and it’s thickness is thought to depend on the interplay between different parameters (such as nutrient concentration, maximal nutrient consumption rate and cell density)^33,35^. Nadell *et. al*. have argued that the (average) thickness of the active layer plays an important role in controlling biofilm spatial structure^33,35^. In this paper, we argue that a careful examination of the active layer dynamics, rather than the average thickness alone, can provide deeper insights into what is driving the changes in biofilm spatial structure, about the distinctive behaviour of the interface roughness under different parameter regimes, as well as the relationship between the mechanical and nutrient behaviours.

**Figure 1.**
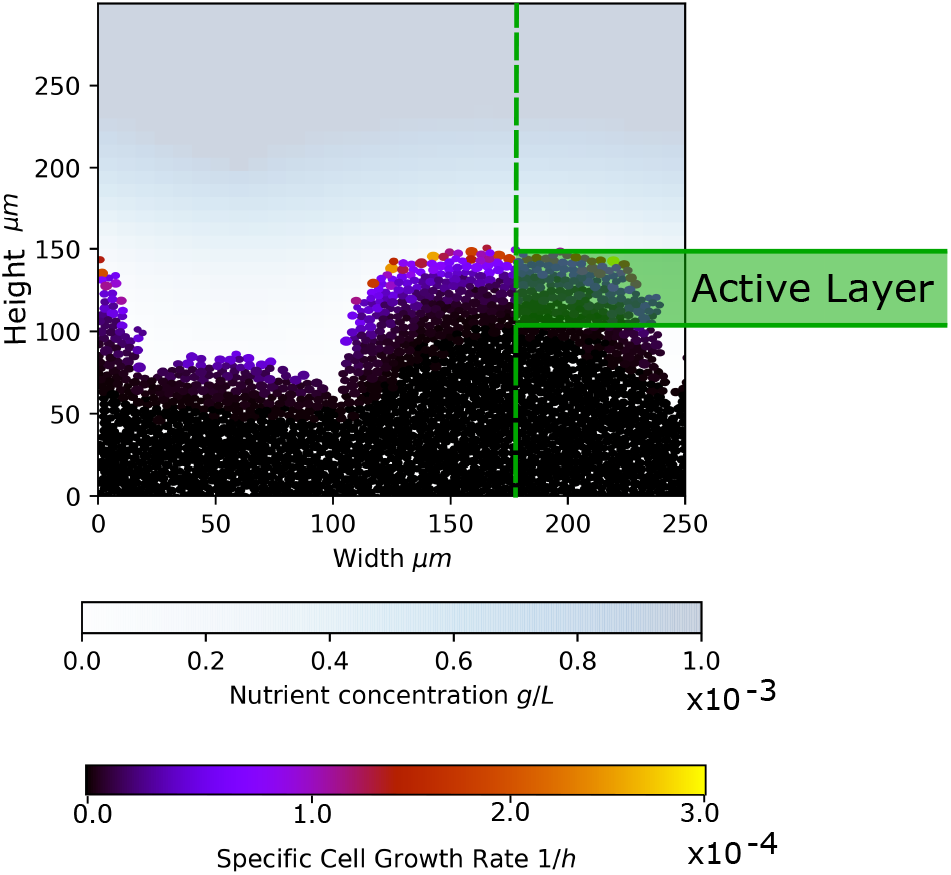
The concept of the active layer. A biofilm configuration generated in our simulations is shown, with the cells in the biofilm colour-coded according to their specific growth rate. The nutrient concentration field is shown on the blue scale. Nutrient is consumed by cells at the top of the biofilm, so that cells deeper in the biofilm are deprived of nutrient and do not proliferate. The active layer is defined as the layer of growing cells at the top of the biofilm (see Methods for further details).

In this paper, we examine the role of active layer dynamics in detail using individual-based simulations of growing biofilms for a range of simulation parameters. We develop a computational method that allows us to simulate biofilm growth over long times, to obtain a clear picture of the steady-state spatial structure. We observe three phases, each with a distinct active layer behaviour and interface roughness trajectory. The first has an unbroken active layer and a smooth steady state roughness (’unpinnned phase’). The second has develops gaps in the active layer that lead to local parts of the interface that become stationary, which we term ‘pinning sites’. These pinning sites are transient in this phase, leading to a fluctuating interface roughness (’transiently pinned phase’). In the third phase, the pining sites remain once they appear, leading to fingering of the biofilm interface and a monotonically increasing interface roughness. To understand what controls the pinning transition in biofilm morphology, we search for a universal ‘control parameter’ that can describe the transition. While others have suggested the average active layer thickness^33^, or a parameter that balances nutrient and mechanical effects^17^ as possible control parameters for biofilm structure, we find that a ratio of the average and standard deviation of the active layer thickness provides a good description of the pinning transition. Since the standard deviation reflects fluctuations in the active layer thickness, this further supports our hypothesis that active layer dynamics play a key role in driving the pinning behaviour of the growing biofilm interface.

## Results

### Individual-based simulations produce distinct biofilm morphologies

To investigate biofilm morphology, we performed individual-based simulations of biofilm growth using the well-established iDynoMiCS simulation software^37^. Our simulations model individual bacteria as discs (in 2D) which consume nutrients, grow, divide, and push each other out of the way (see Methods). The nutrient is assumed to diffuse from above, mimicking approximately an experimental flow-cell setup (see Methods)^37^. We performed a grid of simulations, varying systematically the bulk nutrient concentration *S*_*bulk*_ and the maximum specific growth rate of the bacteria *µ*_*max*_. These parameters have previously been implicated in controlling the average active layer thickness and hence biofilm structure^33^. They could in principle be controlled experimentally by changing the nutrient concentration of the fluid medium in a flow cell setup, and the bacterial strain. All other parameters remain fixed in our simulations (see Table 1).

**Table 1.**
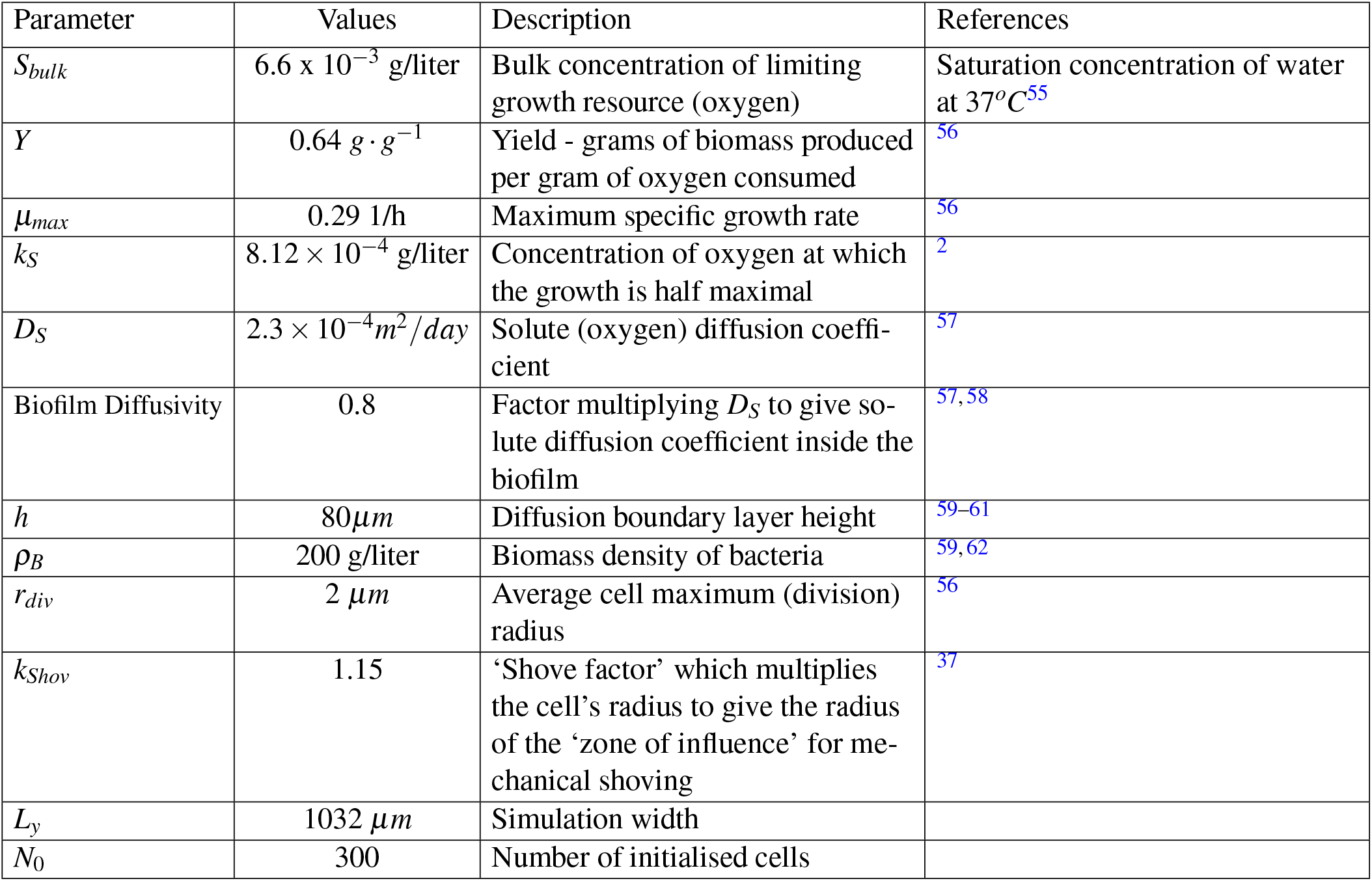
Table of the input values used in our iDynoMiCS biofilm simulations. These values aim to be consistent with *Pseudomonas aeruginosa* in an oxygen-limited flow cell type set up^2,26^.

| Parameter | Values | Description | References |
| --- | --- | --- | --- |
| $S_{bulk}$ | $6.6 \times 10^{-3}$ g/liter | Bulk concentration of limiting growth resource (oxygen) | Saturation concentration of water at 37°C<sup>55</sup> |
| $Y$ | $0.64 \text{ g} \cdot \text{g}^{-1}$ | Yield - grams of biomass produced per gram of oxygen consumed | <sup>56</sup> |
| $\mu_{max}$ | 0.29 1/h | Maximum specific growth rate | <sup>56</sup> |
| $k_S$ | $8.12 \times 10^{-4}$ g/liter | Concentration of oxygen at which the growth is half maximal | <sup>2</sup> |
| $D_S$ | $2.3 \times 10^{-4} \text{ m}^2/\text{day}$ | Solute (oxygen) diffusion coefficient | <sup>57</sup> |
| Biofilm Diffusivity | 0.8 | Factor multiplying $D_S$ to give solute diffusion coefficient inside the biofilm | <sup>57,58</sup> |
| $h$ | 80 $\mu\text{m}$ | Diffusion boundary layer height | <sup>59–61</sup> |
| $\rho_B$ | 200 g/liter | Biomass density of bacteria | <sup>59,62</sup> |
| $r_{div}$ | 2 $\mu\text{m}$ | Average cell maximum (division) radius | <sup>56</sup> |
| $k_{Shov}$ | 1.15 | ‘Shove factor’ which multiplies the cell’s radius to give the radius of the ‘zone of influence’ for mechanical shoving | <sup>37</sup> |
| $L_y$ | 1032 $\mu\text{m}$ | Simulation width | |
| $N_0$ | 300 | Number of initialised cells | |

In our simulations, we observe that the overall biofilm growth rate (cell number vs time) depends strongly on the parameters, with small values of *S*_*bulk*_, or large values of *µ*_*max*_, leading to slow growth (Figure S1). Therefore, to make a fair comparison between biofilms at a similar developmental stage, we chose to compare simulated biofilms of the same size (cell number), rather than age (time). This approach has been taken by others such as Drescher *et. al*., who argue that cell number, rather than time, is the key parameter that determines biofilm architecture^38^.

Figure 2 shows snapshots from our grid of simulations, for biofilms of 75,000 cells. A variety of different biofilm morphologies arise in the simulations, from approximately flat (high *S*_*bulk*_, low *µ*_*max*_) to fingered (low *S*_*bulk*_, high *µ*_*max*_). The growth rate of individual cells within the biofilm is indicated by the colour scale. As expected, our simulations reveal the emergence of an active layer (Figure 1) in which growing cells are localised at the biofilm interface. Figure 3(a) shows the average active layer thickness across the biofilm width as a function of cell number for the simulations in Figure 2 (see Methods for the definition of active layer thickness used here). After an initial period corresponding to biofilm formation, the average active layer thickness reaches a steady state, consistent with previous work^8,32,33,35^. We also quantify the variation in the local active layer thickness using the standard deviation of the active layer thickness, calculated using bins along the biofilm width, where we include empty bins that indicate parts of the interface with no active layer (see the Methods for further details). This is plotted in Figure 3(b), where see that the standard deviation of the active layer thickness takes longer to reach its steady state than the average active layer thickness; highlighting the need for long-time simulations such as those performed here.

**Figure 2.**
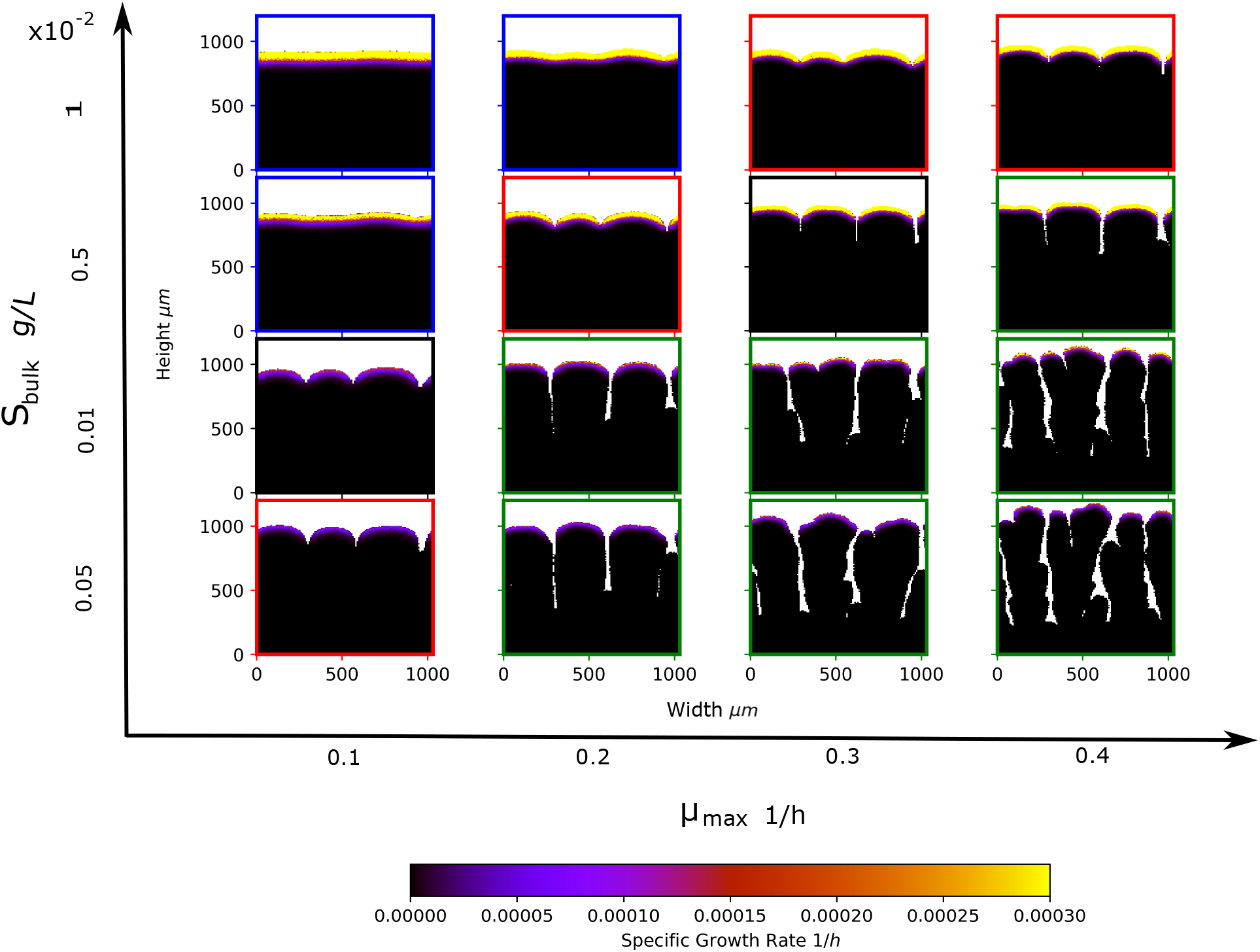
Emergence of distinct biofilm morphologies. Snapshots from our grid of simulations, for biofilm sizes of approximately 75,000 cells. Our grid of simulations was defined by varying the bulk nutrient concentration *S*_*bulk*_ and maximum specific cell growth rate *µ*_*max*_. The remainder of the simulation input parameters are held constant, and detailed in Table 1. In the snapshots, cells are colour coded according to their specific growth rate (due to competition for nutrients this is always significantly less than *µ*_*max*_). The coloured borders around the snapshots indicate the phase of biofilm growth - blue for unpinned, red for transiently pinned, green for pinned and black for transitional, as defined later in Section ‘Biofilm Dynamics show Interface Pinning’ and in Figures 4 and 6.

**Figure 3.**
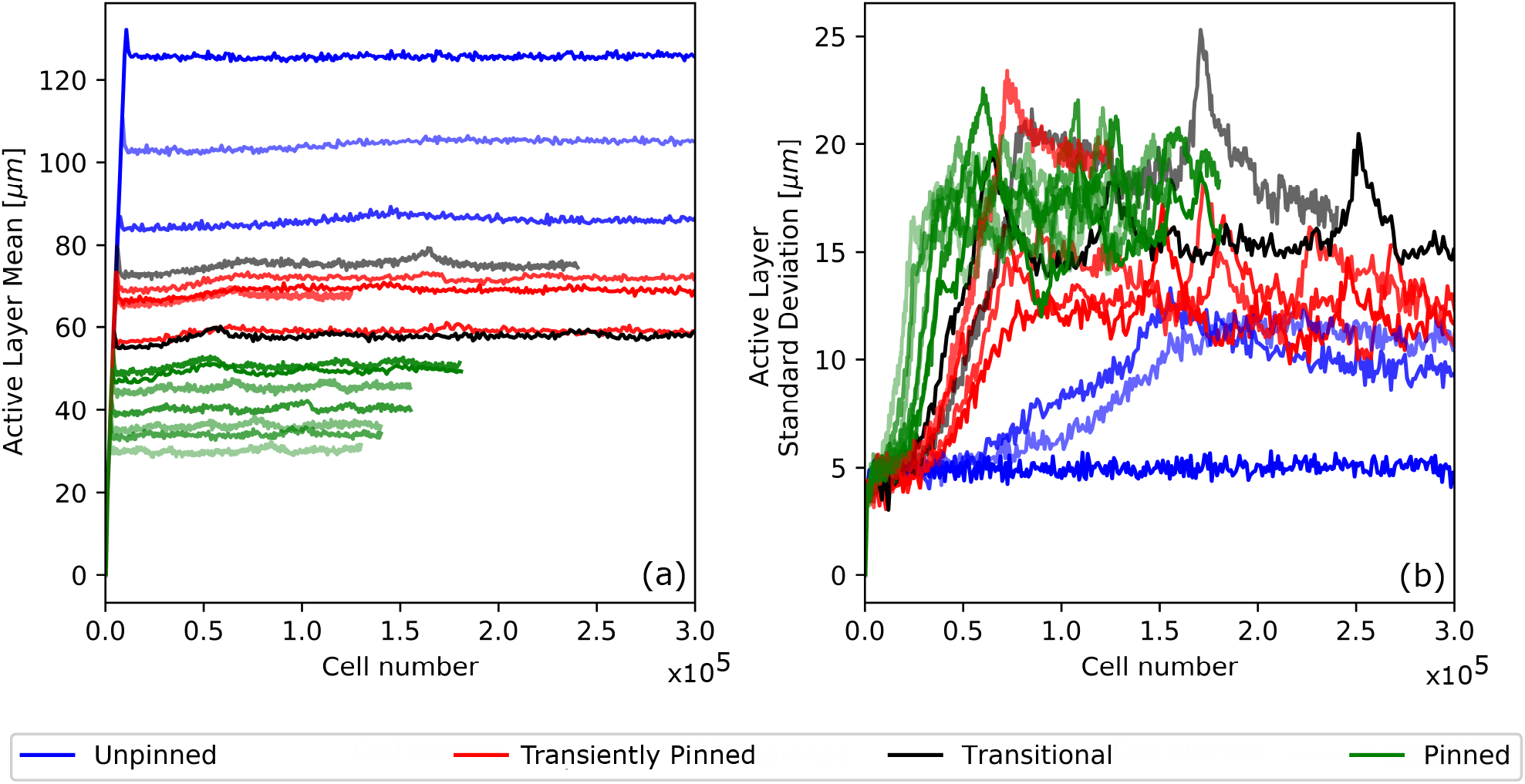
Trajectories of the active layer thickness. The average (panel (a)) and standard deviation (panel (b)) of the active layer thickness are plotted as a function of cell number, for each of the simulations of Figure 2. Here cell number can be viewed as a proxy for time, with the conversion factor being the biofilm growth rate, which is parameter-dependent (see Supplementary Figure S1). The trajectories are colour coded according to their phase - blue for unpinned, red for transiently pinned, green for pinned and black/grey for transitional, as defined in the text and Figures 4 and 6. Supplementary Figures S2 and S3 show these same trajectories individually.

Our simulations clearly show that both the average and the standard deviation of the active layer thickness correlate with the spatial structure of the biofilm. Consistent with previous work^33^, we observe that the active layer is, on average, thick for the smooth biofilms (high *S*_*bulk*_, low *µ*_*max*_) and thinner for the fingered biofilms (low *S*_*bulk*_, high *µ*_*max*_) in Figure 3(a). We also note that the active layer is unbroken in the smooth biofilms, but broken in the fingered biofilms, where growth only occurs at the tips of the fingers (hence the troughs correspond to gaps in the active layer). This is reflected in the values of the standard deviation of the active layer thickness: variations in the active layer thickness are larger in fingered biofilms (low *S*_*bulk*_, high *µ*_*max*_) than in smooth biofilms (low *S*_*bulk*_, high *µ*_*max*_).

### Biofilm dynamics show interface pinning

Figure 3 shows that the average active layer thickness (across the biofilm interface) remains constant over the course of biofilm growth. However, careful inspection of the trajectories of our simulations reveal that there are large local variations in the active layer thickness. We observe three distinct types of dynamical active layer behaviour, shown in Figure 4 and Supplementary movie files. Figure 4 (top row) shows a biofilm that retains a thick, unbroken active layer, with a smooth interface that undulates due to fluctuations in the active layer growth dynamics. Figure 4 (middle row) shows a biofilm in which fluctuations in the active layer become large enough for gaps to form, i.e. regions along the interface where there are no growing cells (e.g. point A). These gaps can cause local regions of the interface to ‘pin’, i.e. to remain stationary as the rest of the growing interface continues to advance (e.g. point B). We call these locally pinned parts of the interface ‘pinning sites’, defining a pinning site as a region of the interface which has not moved over 6 hours simulated time (see Methods). These pinning sites are indicated in red in Figure 4. In the simulation shown in the middle row of Figure 4, the pinning sites are transient, such that local regions of the interface become pinned and later unpinned (e.g. point C). However, in the simulation shown in Figure 4 (bottom row), pinning sites appear and remain fixed. In other words, the interface pins but does not unpin; leading to fingering of the biofilm. Secondary pinning sites may also appear within existing biofilm fingers, causing them to branch. Thus we can define three distinct classes of dynamical behaviour; we refer to these three classes as ‘phases’ of biofilm growth. By analogy with interface growth theory in statistical physics^31^, we denote these the ‘unpinned’, ‘transiently pinned’ and ‘pinned’ phases respectively.

**Figure 4.**
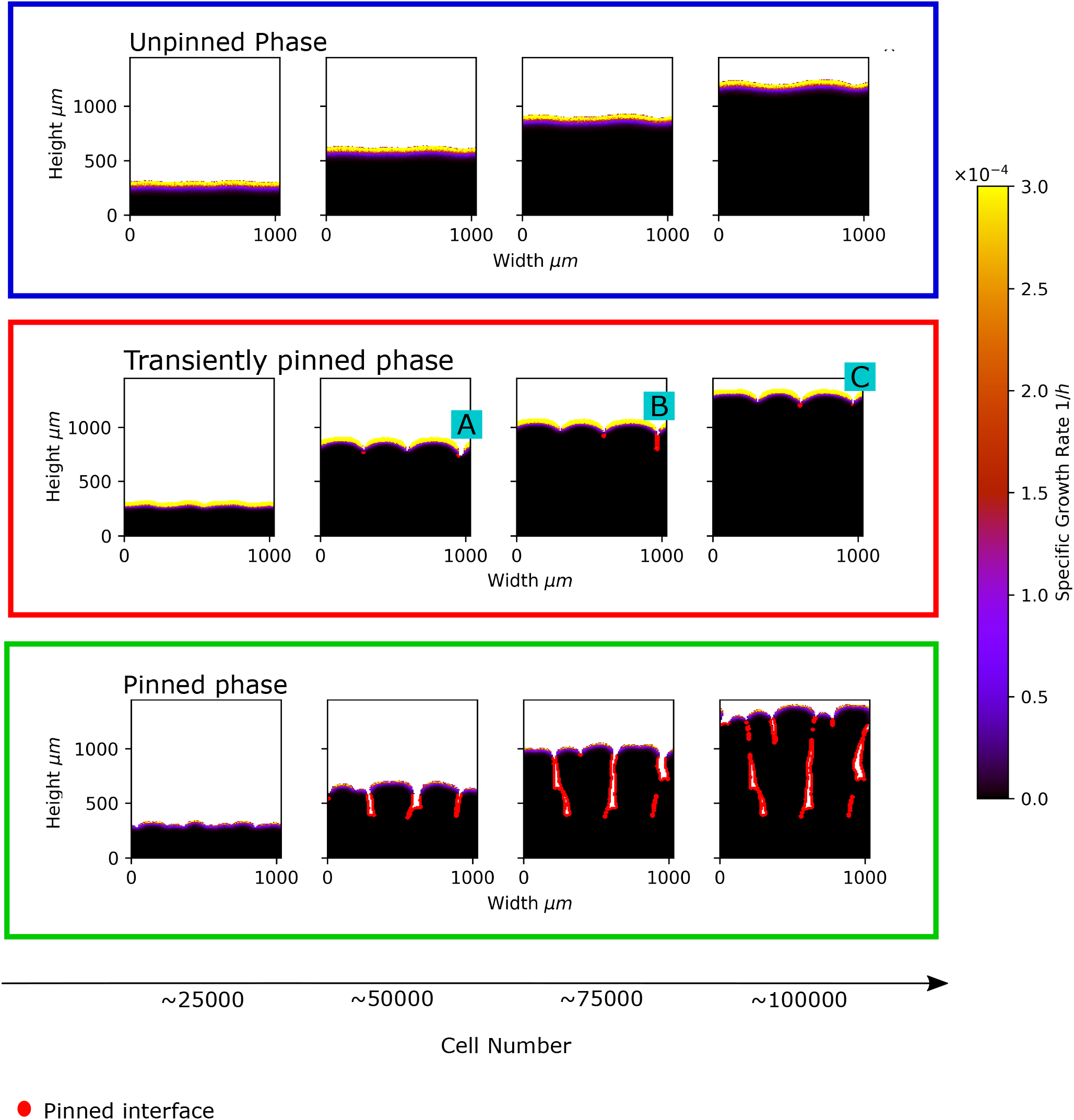
Distinct phases of biofilm dynamics. Biofilm growth is illustrated for three parameter sets, representing three qualitatively different types of biofilm behaviour, or ‘phases’. The top row of snapshots is for parameters *S*_*bulk*_ = 0.01 g/L, *µ*_*max*_ = 0.1 1/h which represent the unpinned phase. The central row of snapshots is for parameters *S*_*bulk*_ = 0.01 g/L, *µ*_*max*_ = 0.4 1/h, representing the transiently pinned phase. The bottom row of snapshots is for parameters *S*_*bulk*_ = 0.0005 g/L, *µ*_*max*_ = 0.4 1/h and represents the pinned phase. Biofilm growth is shown from left to right; snapshots are shown for biofilm sizes of 25,000, 50,000, 75,000 and 100,0000 cells. In the snapshots, cells are colour coded according to their specific growth rate. Parts of the interface that are pinned (i.e. have not moved in the previous six hours of simulated time) are represented in red. The parts of the interface labelled A, B and C in the figure correspond respectively to a gap in the active layer, a pinning site and a gap in the active layer gap after a pinning site has closed.

### Active layer gap dynamics can lead to interface pinning

We can further understand the pinning behaviour in our simulations by looking in more detail at the dynamics of the active layer along the biofilm interface. Figure 5 (left panel) shows an ‘active layer kymograph’ plot for a simulation in the transiently pinned phase (*S*_*bulk*_ = 0.005 g/L, *µ*_*max*_ = 0.2 1/h). The local active layer thickness along the biofilm width (horizontal axis) is shown by a colour scale, such that darker colours represent regions with thin or no active layer and lighter colours represent regions with a thicker active layer. The vertical axis represents biofilm size, revealing the dynamics of the active layer as biofilm growth progresses. In the kymograph, gaps in the active layer are easily visible as dark lines – providing a convenient way to visualise the creation, annihilation, and motion of active layer gaps. The kymograph reveals that active layer gaps appear spontaneously (emergence of a new dark line reading bottom to top in the kymograph) but they disappear only by merging with other active layer gaps. We also observe motion of active layer gaps (sloping of the dark lines in the kymograph), due to pushing between adjacent bulges in the interface (since active layer gaps correspond to the troughs between bulges). This motion is clearly not diffusive (Figure 5; see also Supplementary Figure S4 for the kymographs of our complete grid of simulations).

**Figure 5.**
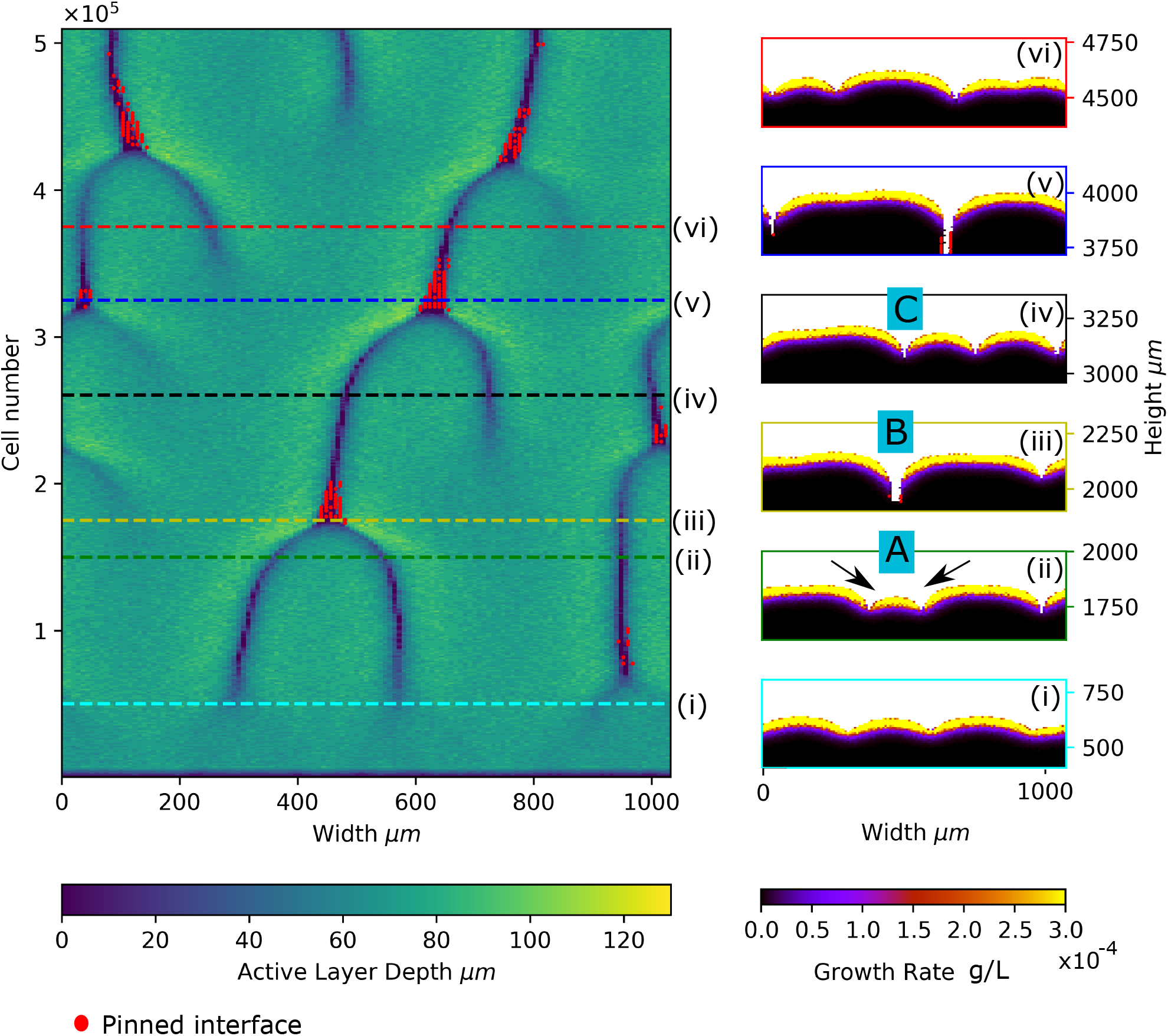
Active layer gap dynamics and pinning. The left panel shows an ‘active layer kymograph’ plot for a simulation in the transiently pinned phase (*S*_*bulk*_ = 0.005 g/L, *µ*_*max*_ = 0.2 1/h). The local active layer thickness (shown by the colour scale) is plotted along the biofilm width (horizontal axis) as biofilm growth progresses (cell number is on the vertical axis). The red dots indicate local regions of the interface that are pinned. Snapshots (i)-(vi) on the right hand side correspond to the dashed lines on the kymograph. Labels A, B and C indicate regions of the interface corresponding to just before, during and after formation of a pinning site.

Figure 5 allows us to connect the dynamics of the active layer to interface pinning. Parts of the interface which are pinned are indicated with red dots superposed on the kymograph. In this simulation, frequent formation and annihilation of pinning sites occurs. The kymograph shows clearly that new pinning sites are formed when two active layer gaps merge. To understand this better, we inspect simulation snapshots corresponding to the kymograph (right panel; the snapshots correspond to the dashed lines in the kymograph). The snapshots show that the merger of two active layer gaps occurs when a small bulge in the interface (labelled A in Figure 5; the arrows indicate the small bulge) is engulfed by the lateral expansion of surrounding, larger, bulges (labelled B and C). We also note that in the simulation of Figure 5, the ‘bulge engulfment’ event that leads to active layer gap merging and formation of a new pinning site is accompanied by the appearance of a new active layer gap at a different point along the interface. Supplementary Figure S4 confirms that this behaviour is common in the transiently pinned phase.

These observations hint that both nutrient consumption and mechanical interactions play a role in the creation of pinning sites in our simulations. Once a bulge in the growing front emerges, it creates an inhomogeneity in the nutrient field by depleting nutrients around it; this leads to gaps in the active layer at the sides of the bulge. Mechanical interactions among the growing cells at the interface lead to pushing between adjacent bulges, which causes the active layer gaps to move. Competition between adjacent bulges leads to the bulge engulfment process that creates new pinning sites in the transiently pinned phase (Labels A-C in Figure 5).

In contrast, the annihilation of pinning sites (which also occurs in the transiently pinned phase) appears to be primarily driven by mechanical interactions. Annihilation of a pinning site happens when a trough in the interface closes up. Careful inspection of movies from our simulations (see Supplementary movies) shows that this occurs because of lateral expansion, i.e. the trough becomes narrower from the sides. This lateral expansion appears to be driven by mechanical pushing interactions that are transmitted from the growing cells at the biofilm front down to the non-growing cells in the walls of the trough. However, the representation of mechanical interactions in our iDynoMiCS simulations is rather crude (see Methods); deeper insight into the roles of nutrient-mediated and mechanical interactions might be gained using simulations that model the pushing interactions between bacteria in more detail^17, 23, 39, 40^.

### Biofilm roughness correlates with pinning dynamics

We now consider the implications of interface pinning for the roughness of the biofilm. The roughness is defined as the standard deviation of the height of the biofilm interface. In our simulations, especially in the pinned phase, we often see interface overhangs, in which the boundary of the biofilm has multiple values for a given position along the horizontal direction (see, e.g. the bottom right panels of Figure 2). To account for this, we define a multi-valued interface and calculate the standard deviation of the vertical position of all points along this interface (see Methods).

Figure 6(a) shows how the roughness of the interface changes as the biofilm grows, in each of our simulations. The trajectories are colour-coded: simulations that correspond to the unpinned, transiently pinned and pinned phases are shown in blue, red and green respectively. We observe qualitatively different roughness trajectories for the three phases of biofilm growth. For simulations in the unpinned phase (blue), the roughness reaches a steady state at a low value, i.e. the interface is rather smooth. For simulations the transiently pinned phase (red), the roughness reaches a steady state with large fluctuations. For simulations in the pinned phase (green), the roughness does not reach a steady state, but instead increases monotonically throughout the simulation. We also see some simulation trajectories which do not fit easily into these categories; these are shown in black and grey in Figure 6. These simulated biofilms may be undergoing transitions between the different phases of biofilm growth; for example the trajectory shown in grey appears to start in the transiently pinned phase and transition to the pinned phase.

**Figure 6.**
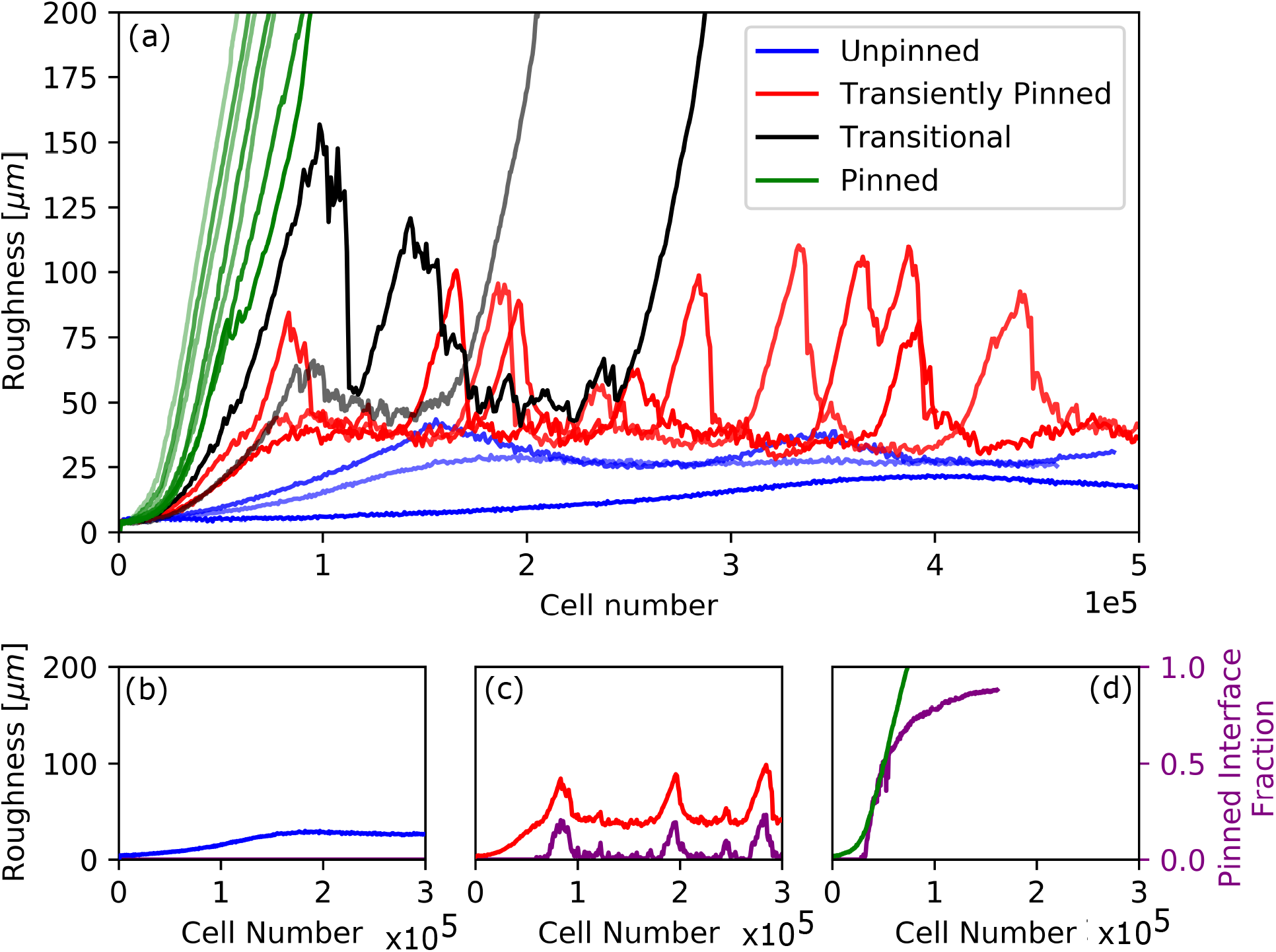
Biofilm roughness dynamics driven by interface pinning. Panel (a) shows roughness trajectories (roughness vs cell number) for each of the simulations of Figure 2, i.e. for varying *S*_*bulk*_ and *µ*_*max*_. Trajectories in blue, red and green correspond to simulations in the unpinned, transiently pinned and pinned phases respectively, as defined by their interface pinning behaviour (see Figure 4). The trajectories shown in black and grey appear to be transitioning between phases. Panels (b)-(d) show a sample roughness trajectory for each of the phases (blue/red/green line) plotted together with the corresponding trajectory for the fraction of the interface that is pinned, i.e. that has not moved in the previous 6 hours of simulated time (purple line). Panels (b)-(d) show the same simulations for each phase as Figure 4, i.e. *S*_*bulk*_ = 0.01 g/L, *µ*_*max*_ = 0.1 1/h for the unpinned phase, *S*_*bulk*_ = 0.01 g/L, *µ*_*max*_ = 0.4 1/h for the transiently pinned phase and *S*_*bulk*_ = 0.0005 g/L, *µ*_*max*_ = 0.4 1/h for the pinned phase. Supplementary Figures S5 and S6 show these same trajectories of interface roughness and pinned interface fraction for all of our simulations individually.

The distinct behaviour of the interface roughness in the three phases of biofilm growth can be understood in the context of the interface pinning dynamics. To quantify pinning, we measure the fraction of the interface that is pinned i.e. stationary, where we define as the parts of the interface that are pinned as those that do not move for at least 6 hours of simulated time, as in Figure 4 (see Methods). In Figure 6(b)-(d) we superpose trajectories of the interface roughness and the pinned interface fraction for representative simulations from each of the three phases. In the unpinned phase (Figure 6(b)) the interface is not pinned and the interface is rather smooth. The transiently pinned phase (Figure 6(c)) is characterised by transient pinning of the interface, such that the pinned fraction of the interface undergoes large fluctuations. Figure 6(c) shows that these correlate directly with the observed fluctuations in interface roughness. This makes intuitive sense, since pinned regions of the interface (by definition) remain stationary while the rest of the interface continues to advance. Therefore the width of the interface (the difference between maximum height and minimum height) increases while a pinning site is present. Once the pinning site is annihilated, the trough in the interface closes up and the interface roughness decreases. Figure 6(d) shows the equivalent plot for the pinned phase. Here, pinning sites appear and persist, leading to fingering of the interface. This corresponds to a pinned interface fraction that increases monotonically, tending towards unity. Since the biofilm fingers continue to grow throughout the simulation, the width of the interface, and hence the roughness, also increases throughout the simulation. In a similar manner, we can link the behaviour of the interface roughness (i.e. the interface fluctuations) to the behaviour of the standard deviation of the active layer thickness (i.e. the active layer fluctuations); see Supplementary Figure S7.

### Towards a phase diagram for biofilm pinning

Our work so far suggests that a growing biofilm can undergo a transition from a state with an unbroken active layer and a smooth interface, to a state in which the active layer has gaps, with local parts of the interface being pinned, and a rough interface. The latter state may be ‘transiently pinned’ (with short-lived pinning sites) or ‘pinned’ (with permanent pinning sites). To understand better the nature of this transition, we attempt to plot a phase diagram. A phase diagram is a central concept in the physical sciences, used to describe how a system transitions from one state to another. An order parameter describes the physical state of the system, and a control parameter describes the environment; the phase diagram shows the order parameter plotted as a function of the control parameter^41,42 1.^ The nature of the order parameter and control parameter, and the shape of the resulting phase diagram, can reveal important information about the key physical principles underlying the transition.

To construct a phase diagram for interface pinning in our simulations, we therefore aim to plot an order parameter, describing the extent of interface pinning, as a function of a control parameter. Since the distinguishing feature of the three phases of biofilm growth is the presence of pinned regions of the interface, we choose as our order parameter the average steady-state fraction of the interface that is pinned (see Methods). As shown in Figure 6, this is a well-defined quantity that takes different values in the three phases of biofilm growth. In the unpinned phase it is zero, while it lies in the ranges 0.010-0.284 and 0.741-0.875 for our simulations that have previously been identified as being in the transiently pinned and pinned phases respectively.

We now search for the best control parameter to describe the biofilm pinning transition. We define a control parameter as optimal if it causes our simulation data to collapse onto a single curve in the control parameter - order parameter space. In previous work, it has been suggested that the average active layer thickness drives biofilm spatial structure, with a thick active layer leading to smooth biofilms while a thin active layer leading to rough biofilms^33,35^. Therefore we begin by considering the steady-state value of the average active layer thickness (Figure 3(a)) as a candidate control parameter. Figure 7(a) shows the order parameter (steady-state pinned interface fraction) plotted as a function of the average active layer thickness. Here we include all the simulations of Figure 4 (apart from the transitional simulations and one which has not reached the steady state), together with additional simulations in the transiently pinned phase to gain resolution in the transition region. The data points are colour coded according to phase of biofilm growth; simulations in the unpinned, transiently pinned and pinned phases are shown as blue, red and green data points respectively. Figure 7(a) shows that, as candidate control parameter, the average active layer thickness does distinguish between the three phases, but it does not produce a perfect collapse of the data onto a single line, particularly in the transiently pinned phase. This suggests that the average active layer thickness is not the only factor controlling the pinning transition.

**Figure 7.**
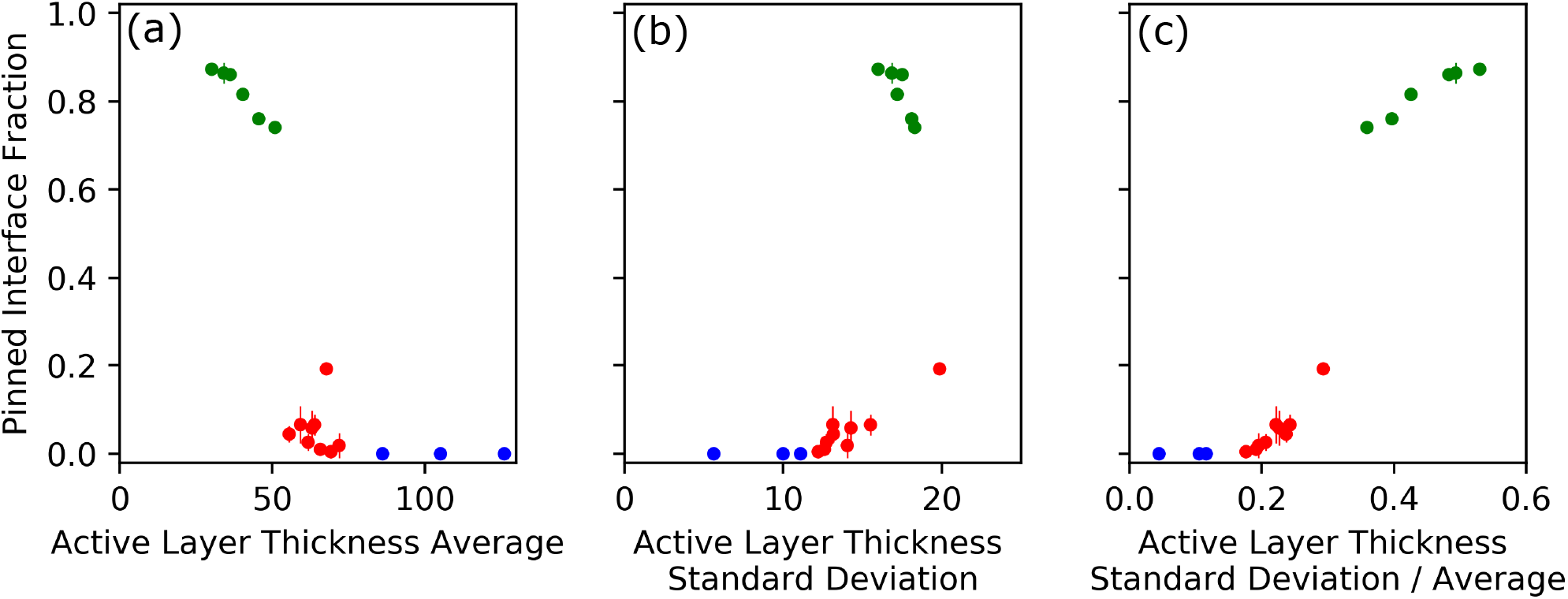
Phase diagram for the pinning transition. In each panel, our chosen order parameter, the average steady state pinned interface fraction, is plotted against three potential control parameters for the data from our simulations. These are the average active layer thickness (panel (a)), the steady state of the standard deviation of the active layer thickness (panel (b)), and the steady state coefficient of variation (i.e. standard deviation / mean) of the active layer thickness (panel (c)). Simulation data points are coloured according to whether the simulation is in the unpinned (blue), transiently pinned (red) or pinned (blue) phase. The simulations included in our plot are as in Figure 4, but the transitional simulations (see Figure 6) are not included. The simulation with *µ*_*max*_ = 0.4 1/h,*S*_*bulk*_ = 0.005 is also not included, as it had not fully reached the steady state (see Supplementary Figure S6). To gain resolution in the transition region, we also included additional simulations in the transiently pinned phase, with parameter values of *µ*_*max*_ = 0.3 1/h,*S*_*bulk*_ = 0.007 g/L;*µ*_*max*_ = 0.33 1/h,*S*_*bulk*_ = 0.01 g/L, *µ*_*max*_ = 0.45 1/h,*S*_*bulk*_ = 0.01 g/L; *µ*_*max*_ = 0.25 1/h,*S*_*bulk*_ = 0.005 g/L; *µ*_*max*_ = 0.37 1/h,*S*_*bulk*_ = 0.01 g/L. Error bars are calculated using the standard error of the mean for correlated data (see Methods).

Motivated by our previous observations about active layer dynamics (Figure 4 and Supplementary Figure S4) we decided to investigate whether *fluctuations* in the active layer might also be a driver for the pinning transition. The standard deviation of the active layer thickness, measured across the biofilm width, provides a crude measure of active layer fluctuations, and it reaches a well-defined steady state in our simulations (Figure 3; see also Supplementary Figure S7 where the behaviour of the standard deviation of the active layer thickness is contrasted with that of the interface roughness). Therefore we chose the steady-state value of the standard deviation of the active layer thickness as a second candidate control parameter. Figure 7(b) shows the resulting candidate phase diagram. This candidate control parameter also distinguishes between the three phases, but the transiently pinned phase data (red) does not collapse perfectly, and the pinned phase data (green) shows a strange behaviour where the order parameter decreases with increasing control parameter.

Finally, we reasoned that perhaps the coefficient of variation of the active layer thickness, i.e. the standard deviation divided by the mean, might make a better control parameter. This makes intuitive sense, since we expect that the creation of an active layer gap (which can lead to a pinning site) requires the active layer thickness to fluctuate by an amount that is comparable to its mean value. Figure 3(c) shows our third, and final, candidate phase diagram, in which the control parameter is the coefficient of variation of the active layer thickness. This produces a better data collapse than our previous candidate control parameters (Figure 3(a) and (b)). As the coefficient of variation of the active layer thickness increases, the system transitions from the unpinned to the transiently pinned to the pinned phase. Therefore we conclude that (i) active layer fluctuations are important in driving the biofilm pinning transition and (ii) the magnitude of these fluctuations relative to the mean active layer thickness is the relevant quantity.

The form of the phase diagram can also tell us about the fundamental physics of the system. In general terms, we can distinguish a ‘discontinuous transition’, in which the order parameter jumps discontinuously from zero to a finite value at a critical value of the control parameter, from a ‘continuous transition’, in which the order parameter changes continuously from zero to a finite value as the control parameter varies (see sketch in Supplementary Figure S8)^41^. In practice, the distinction between discontinuous and continuous transitions can become blurred by finite size effects (one only observes a true discontinuity in the phase diagram for systems of infinite size)^43^. For our system, in Figure 3(c), the variation in the order parameter with the control parameter appears at first sight continuous, but there is a large jump in the value of the order parameter between the transiently pinned and pinned phases. Therefore, we tentatively suggest that this may be a discontinuous transition, with the apparent smoothing arising from finite size effects. In this scenario, the transiently pinned phase would arise only in systems of finite size (including real biofilms), while a hypothetical biofilm of infinite lateral size would transition directly from the unpinned to the pinned state upon varying the control parameter. This might also suggest that the transition to a pinned state is a nucleation phenomenon (see Discussion and Supplementary Figure S9).

We note that the phase diagram of Figure 3(c) does not provide practical information about how the biofilm will react to a change in our simulation parameters, since we do not know how our chosen control parameter depends on the simulation parameters. In future, it may be possible to determine how the coefficient of variation of the active layer thickness depends on the parameters of the simulation, in a similar way to previous work on the average active layer thickness^33^. For the moment, though, this phase diagram provides conceptual rather than practical information.

## Discussion

In this work, we used individual-based computer simulations to investigate the spatial structure of growing bacterial biofilms. Our simulations include the effects of local nutrient limitation (modelled via a reaction-diffusion equation) and mechanical pushing between the cells (modelled crudely via a ‘shoving’ algorithm; see Methods). Varying the nutrient concentration and the maximal specific growth rate of the bacteria we observed a diversity of biofilm morphologies, ranging from smooth to highly fingered interfaces. Our simulations could be classified into three ‘phases’ of biofilm growth, with the key distinguishing feature being the pinning behaviour of the biofilm interface. In the ‘unpinned’ phase the interface is smooth and does not pin, and the active layer is thick and unbroken. The ‘transiently pinned’ phase is characterised by the appearance of transient local pinning sites along the interface; these pinning sites arise because gaps develop in the active layer. The interface roughness shows large temporal fluctuations, which correlate with the transient presence of pinning sites. In the ‘pinned’ phase, the interface develops fingers, which arise from pinning sites that appear but do not disappear. Correspondingly, the interface roughness increases throughout the simulation.

Our results point to a key role for the dynamics of the active layer in driving biofilm spatial structure. The formation of a gap in the active layer is controlled by the magnitude of fluctuations in active layer thickness, relative to the mean thickness; in other words, by the coefficient of variation of the active layer thickness. Once an active layer gap has formed, the dynamics of these local gaps gives rise to pinning sites; for example in the transiently pinned phase, pinning sites emerge when two active layer gaps merge, or in other words when a small bulge in the interface is engulfed by adjacent large bulges. Pinning sites lead to an increase in interface roughness, as the pinned interface remains stationary as the growing front continues to advance. It has already been reported that the average thickness of the active layer correlates with biofilm spatial structure^33^; our work provides the additional insight that active layer *dynamics* is also important, since it drives the emergence of pinning sites.

The relative importance of nutrient limitation and mechanical interactions in controlling biofilm spatial structure has been debated in the literature. Nutrient limitation can drive a fingering instability in the biofilm interface^15^, but recent work suggests that mechanical interactions also drive spatial structure, particularly for smooth biofilms^17,18,23^, as well as in tissue growth more generally^44^. We speculate that in our simulations, creation of pinning sites may be mediated by an interplay between nutrient limitation that drives the growth of interface bulges and hence the creation of gaps in the active layer^15^ and mechanical pushing between adjacent bulges that leads to pinning of the interface; in contrast, pinning site annihilation may be driven primarily by mechanical pushing interactions, generated when growing cells in the active layer push against the non-growing cells that form the walls of a trough in the interface.

To understand better the nature of the biofilm pinning transition, we attempted to construct a phase diagram, i.e. a plot of an order parameter (here the fraction of the interface that is pinned) as a function of a control parameter. In many physical processes a phase transition is controlled by an external parameter (e.g. temperature). Here, however, there is no external parameter that directly links to interface pinning; instead, pinning sites are an emergent feature of the complex process of biofilm growth. In this scenario, choosing the control parameter that produces a good data collapse onto a universal phase diagram can provide fundamental insight into what drives the transition. Motivated by our and others’ findings that the active layer is important in biofilm structure, we tested the average active layer thickness, and the standard deviation of the active layer thickness (across the biofilm width), as candidate control parameters. We found that neither of these produced a very good data collapse, but the coefficient of variation of the active layer thickness (sd/mean), produced a better data collapse. This finding strengthens our view that fluctuations in the active layer are important in controlling biofilm spatial structure.

It would of course be useful to predict biofilm structure given a set of controllable input parameters. To do this, we would need to know how our control parameter, the normalised standard deviation of the active layer thickness, depends on the system parameters. The parameter dependence of the average active layer thickness has been investigated in previous work^33^; it would be interesting (although challenging) to perform a similar analysis for the standard deviation of the active layer thickness. Parameters describing mechanical interactions^17^ as well as those describing the nutrient reaction-diffusion equations, would be important here.

The phase diagram that emerges from our work (Figure 3(c)) leads us to speculate that the biofilm pinning transition is discontinuous in the control parameter, with the transiently pinned phase arising from finite size effects. This distinction has relevance for the kinetics of the phase transition since, for equilibrium systems, a discontinuous transition implies a that stochastic fluctuation is required to overcome an activation barrier (i.e. it is a nucleated process), while for a continuous transition the transition happens spontaneously^41^ (see Supplementary Figure S9). Though the kinetics of phase transitions are less well defined for non-equilibrium systems such as our growing biofilms, our work may hint that a critical fluctuation may be needed to initiate a structural phase transition (see Supplementary Figure S9). From a mechanistic point of view, this could correspond to the appearance of a gap in the active layer that is wide enough that it does not close up again, or a pinning site that is wide enough that it does not depin. Interestingly, if pinning events are indeed activated processes this would contrast with the classic work of Dockery and Clapper on fingering instabilities in biofilm growth, which suggests that, due to the inhomogeneous nutrient field, the interface is unstable to perturbations of any size^15^. Dockery and Clapper did not, however, take account of mechanical interactions.

In this work, we have taken care to study biofilm growth over long times, once a steady state has been reached; this required the development of a computationally efficient clipping algorithm (see Methods and Supplementary Material). We note that at earlier times in our simulations, the interface roughness can appear to reach a plateau (even in the pinned phase), before later increasing (Supplementary Figure S10). Therefore, in shorter simulations it may be hard to know whether the true steady state has been reached. Our long-time simulations suggests that, in the pinned phase, the interface roughness does not in fact reach a steady state, but rather continues to increase because the tips of the fingers continue to grow while the interface remains pinned at the troughs. In contrast, a finite steady-state roughness would correspond to an interface that has stalled in its net growth. From a practical point of view, the fact that the biofilm fingers continue to grow in our simulations presents computational issues, since the use of our clipping algorithm is constrained in the case of fingered biofilms (see Methods). This means that while the steady state of the active layer dynamics and interface roughness behaviour can be reached, it remains challenging to reach the full steady state of the pinned interface fraction in the case of the pinned biofilms (see, e.g. Supplementary Figure S6).

Our work has an interesting analogy with pattern formation in crystal growth, where complex crystal morphologies arise from instabilities in the advancing solidification front^45^. The local rate of crystal growth can be limited by the rate of diffusion of heat away from the crystallisation front (e.g. crystallization of small molecules or metals), or by the rate of diffusion of molecules to the growth front (e.g. in polymer crystallization). This leads to fingering instabilities similar to the nutrient-driven fingering instability in biofilm growth^15^. The emergence of different crystal forms might also have parallels with the emergence of mutant clones in a biofilm. However, we note that in crystal growth the morphological instability is ultimately limited by surface tension, which tends to smooth the interface^45^, while the limiting factor for biofilm growth is less clear (although it probably involves mechanical interactions). We also note that in a crystal, growth occurs only right at the interface, while for a biofilm, there is a growing region of finite thickness at the interface (the active layer).

Our work also has a clear connection with the statistical physics of pinning-depinning transitions in interface growth. Here, diverse interface growth phenomena are grouped into a small number of ‘universality classes’, based on their scaling behaviour (the values of the exponents in plots of, e.g. roughness vs time). Phenomenological stochastic differential equations are then used to describe interface growth within a particular universality class; for example the classical Kardar-Parisi-Zhang (KPZ) equation describes fluctuations of an unpinned growing interface, while addition of a quenched noise term to the KPZ equation leads to a model for interface pinning. This ‘quenched KPZ (qKPZ)’ model shows a flat phase with no pinning sites, a pinned phase and a intermediate phase in which pinning sites are overcome^31,46^ (although it is unclear whether this model predicts monotonically increasing roughness in the pinned phase). The qKPZ equation has been applied to biofilm growth^28,29^, but the source of the quenched noise is usually ascribed to external inhomogeneities in the environment. In our simulations, there are no such inhomogeneities; rather, pinning arises from the spontaneous emergence of gaps in the active layer. In the interface growth theory literature, KPZ-type models also exist where interface pinning arises from internal fluctuations in the growth process^46–48^, or where the growing interface is coupled to a nonequilibrium field (such as a nutrient field)^49^. However the relevance of such models for bacterial biofilms and colonies has not been investigated. It is also possible that biofilm growth might be described by an alternative type of interface growth model, such as Diffusion Limited Aggregation^50^. While our focus here was on a more mechanistic analysis of the role of the active layer, it would certainly be interesting in future work to clarify the connection with interface growth theory by measuring the scaling exponents for biofilm growth in individual-based simulations.

From a biological point of view, our simulations are, of course, far from realistic. Perhaps most importantly, our model does not include the extracellular matrix (EPS), which means that our simulated biofilms have a much higher cell density than flow cell biofilms formed by, e.g. *Pseudomonas aeruginosa*. We would also expect the mechanical properties of the biofilm to be strongly influenced by EPS^19^. This could affect the predictions of the model, since, for example, EPS-mediated interactions might act non-locally (between cells relatively far apart in the biofilm), which could alter the phase behaviour. In contrast, bacterial colonies formed by lab strains of *Escherichia coli* produce little or no EPS, so our findings might be more realistic in that context. We also neglect many other features of real biofilms such as fluid flow, chemical signalling between cells, and phenotypic changes associated with biofilm growth. The size of our simulated biofilms is also highly unrealistic. In order to reach the steady state, which is necessary for a rigorous analysis of the underlying physics, our simulations generate extremely thick biofilms, much thicker than those seen in experimental flow cell experiments.

From a physical and/or computational point of view, our work also has limitations. In particular, our simulations are performed in two dimensions, for reasons of computational feasibility. The dimensionality of a system can have profound effects on phase transitions^31^: therefore it will be important to determine in future work whether the same phenomena occur in 3D models. Our interpretation of the results, especially of our phase diagram, may also be limited by our finite system size. In future, it would be interesting to perform simulations at different system sizes to determine whether the transiently pinned phase can indeed be attributed to a finite size effect. Even if this is the case, the transiently pinned phase may still be of some relevance, since real biofilms are of course finite in size. Finally, we also note that the representation of mechanical interactions in our simulations is very crude (the algorithm simply resolves overlaps due to growth by a random ‘shoving’ algorithm; see Methods and Ref.^37^). Other algorithms for simulating bacterial growth represent physical interactions more realistically^17,39,51,52^, and it would be interesting to extend this work in future using such algorithms.

In spite of these caveats, our simulations reveal fundamental insights into the spatial structure of growing biofilms. Specifically, our work reveals a key role for the dynamics of the active layer in driving the creation and annihilation of pinning sites at the biofilm interface, resulting in transitions in spatial structure, with drastic effects on the interface roughness.

## Methods

### Simulation Methods

In this work, we use individual-based biofilm modelling software iDynoMiCS^37^. Briefly, iDynoMiCS models the bacteria in a biofilm as individual agents whose behaviour is coupled to a solute reaction-diffusion equation^37^. The agents, which are assumed to be discs in continuum 2D space, each grow with specific growth rate *µ* according to the Monod equation:

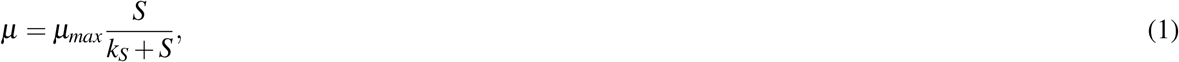

where *µ*_*max*_ is the maximum specific growth rate of the bacteria, *k*_*S*_ is the concentration of the solute at which the growth is half maximal, and *S* is the local solute concentration of the bacterial cell^53^. Once a bacterial cell reaches a maximum radius (which has a stochastic element), it divides into two daughters. Bacteria interact with one another mechanically via a shoving algorithm. Briefly, this algorithm detects pairs of bacteria whose ‘zones of influence’ (defined to be the radius multipled by a ‘shove parameter’) overlap, and shuffles bacterial positions to avoid such overlaps^37^. Although iDynoMiCS has the facility to model extra-cellular matrix (EPS) as non-replicating particles, we did not model EPS in this study. In iDynoMiCS, the solute is represented by a concentration field which varies in space and time due to to diffusion and consumption by the bacteria. A separation of timescales is assumed, such that the reaction-diffusion equation for the solute is assumed to be reach steady state faster than the timescale for bacterial growth; hence the solute concentration equations are solved to steady state for each interaction of the bacterial growth updates. The computational domain is set up to resemble a flow cell, where the biofilm grows on a hard surface and nutrients diffuse from above. Convective flow is not modelled, but rather it is assumed that there is a stationary layer of fluid close to the biofilm (the ‘boundary layer’)^37,54^. It is also assumed that the diffusion constant for solute is reduced inside the biofilm by a fixed factor compared to outside the biofilm. The input values we use in our simulations are based on experimental values for oxygen-limited *Pseudomonas aeruginosa* biofilms, as outlined in Table 1.

In order to reach long simulation time scales, we also use an additional ‘clipping’ algorithm in combination with iDynoMiCS. This algorithm periodically removes inactive cells far below the growing front, such that a computationally feasible number of cells remain in the simulation space. This is achieved by pausing the iDynoMiCS simulation and removing the relevant cells, or ‘clipping’, and then restarting the simulation. This clipping procedure is done at regular time intervals, such that each complete biofilm simulation consists of *N* segments, each of length (in time) *T*_*s*_, producing a total simulated time *T* = *NT*_*s*_. In the clipping procedure, bacterial cells which are both below the lowest actively growing cell and below the minimum point of the interface (which can be different points depending on the biofilm configuration), are removed. The complete algorithm is shown in the Supplementary Information. We rigorously tested this algorithm to ensure it does not perturb biofilm growth (see Supplementary Figures S11 and S12).

### Characterisation of Spatial Structure

As we have seen, our analysis focuses on the active layer, which we define as the layer of growing cells at the top of the biofilm. More specifically, we begin by defining a threshold growth rate; cells which grow faster than this rate are defined to be part of the active layer. We consider a cell to be in the active layer when it grows at greater than 0.1% of the maximal specific growth rate *µ* = *µ*_*max*_*S*_*bulk*_*/*(*k*_*S*_ + *S*_*bulk*_) that is possible under the conditions of the simulation (ie. for given values of *µ*_*max*_ and *S*_*bulk*_). Therefore the condition for a cell to be part of the active layer is

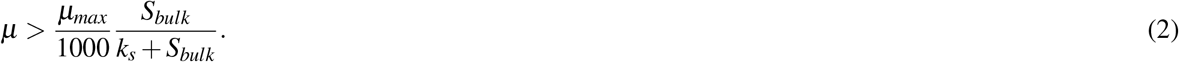

We now outline how the average and standard deviation of the active layer thickness are calculated. We define a grid spanning the simulation domain with *D* columns (horizontal bins) and *H* rows (vertical bins) of width 8*µm*. For each of the *D* columns, we find the total number of ‘active’ grid squares whose biomass has an average specific growth rate above the threshold in Equation 2. This defines the local active layer thickness. For some biofilm configurations, for example if the biofilm is rough, there can be a growing layer both at the leading edge of the biofilm and within a trough, so the active grid squares are not necessarily adjacent to one another. Once the active layer width for each vertical strip has been found, the mean active layer thickness is found by averaging over all the *D* columns. The standard deviation of the active layer thickness is in turn calculated as the standard deviation of the active layer thickness of each of the columns i.e. it is the standard deviation of the local thickness of the active layer for a particular configuration. An active layer gap occurs when a vertical column on the grid contains no active grid squares.

To calculate the interface roughness, we first define the interface boundary. We use a multi-valued interface^63,64^ to correctly account for interface overhangs. On our grid of *D* 8*µm* columns and *H* 8*µm* rows, we search for grid squares which contain biomass but have a nearest neighbour that does not contain biomass. This produces a set of grid squares corresponding to the interface {*k}, k* = 1, …*N*_*int*_ where *N*_*int*_ ≥ *D*. Defining the interface in this way allows there to be multiple vertical points for each horizontal point along the boundary - hence its description as a multi-valued interface.

We then define the interface width, i.e. the interface roughness, as the root mean square height of the points on the interface:

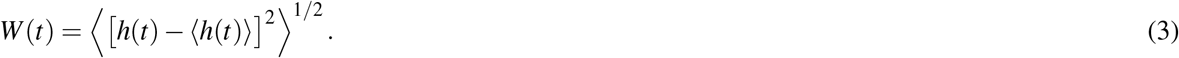

Here, ⟨*h*⟩ is defined as the mean value of *h*_*k*_, where *h*_*k*_ is the vertical coordinate of the *k*th point along the interface, namely,

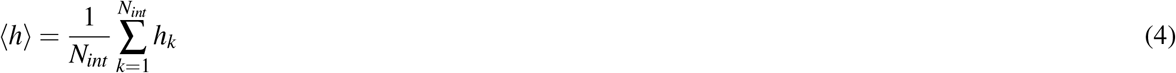

with *N*_*int*_ being the number of points along the interface.

We also define the stationary, or pinned, interface fraction. Here, we define the interface boundary as above, then look for parts of the interface which are both inactive and have not moved in a six hour period (this being the frequency of output files). Specifically, we compare the positions of those interface grid squares which are inactive (i.e. they do not meet the condition of Eq. (2), with the interface boundary of the configuration 6h earlier. If this interface point is common to both configurations, it is defined to be both inactive and pinned. The pinned interface fraction *f*_*P*_ is then defined as

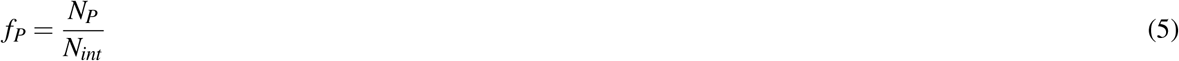

where *N*_*P*_ is the number of inactive, pinned, interface points and *N*_*int*_ is the number of points on the interface.

The error bars in Figures 7 (a)-(c) are calculated as the standard error of the mean (*SEM*) of the *n* time points that were recorded for each variable (average active layer thickness, standard deviation of the active layer thickness and the pinned interface fraction) once the steady state had been reached, adjusted for the fact that our time series are correlated data. The auto-correlation time *τ* was calculated using code from Martin *et. al*.^65^. The effective number of independent data points *n*_*e f f*_ in our correlated data could be calculated as *n*_*e f f*_ = *n/τ*. Finally, the standard error of the mean (*SEM*) was calculated as 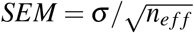, where *σ* is the standard deviation of the *n* steady-state time points.

## Acknowledgements

The authors thank Chris Brackley, Martin Evans, Rhoda Hawkins, Jamie Hobbs, Erik Maikranz and Bartek Waclaw for useful discussions, and David McKain and Donald Grigor for computing support. This work was supported by the European Research Council under Consolidator grant 682237 EVOSTRUC, by the Engineering and Physical Sciences Research Council via a DTA studentship to EY and by the Biotechnology and Biological Sciences Research Council under grant BB/R012415/1.

## Footnotes

1 For example, in the classic textbook example, the order parameter is the degree of magnetisation of a magnetic material while the control parameter is its temperature.

