## Supplemental Material for "Pinning transition in biofilm structure driven by active layer dynamics"

---

**Algorithm 1 The Clipping Procedure Algorithm.** This procedure is used in combination with the iDynoMiCS software to reach long simulation times. This involves running many short simulations with the output biofilm configuration of one simulation segment being clipped and then used as the starting configuration of the next simulation segment. The conditions for determining the threshold height are based on the position of growing cells and on the position of the interface, to avoid perturbing the active layer or the biofilm interface.

---

1. Run the iDynoMiCS simulation from the starting configuration up to segment time $T_s$
2. Calculate the threshold height:
   (a) Calculate the minimum interface height and the minimum of the growing layer
   (b) Set the threshold height as the lowest of the minimum interface height and the minimum of the growing layer
   (c) Subtract a buffer of $20\mu m$ from the threshold height
   (d) If the threshold height is greater than 200 $\mu m$, set it to 200 $\mu m$.
3. Remove cells below the threshold height to create the clipped cell configuration
4. Restart the simulation using the clipped cell configuration, run up to time $T_s$
5. Repeat steps 2-5 $N$ times until the end time $T = NT_s$ is reached

---

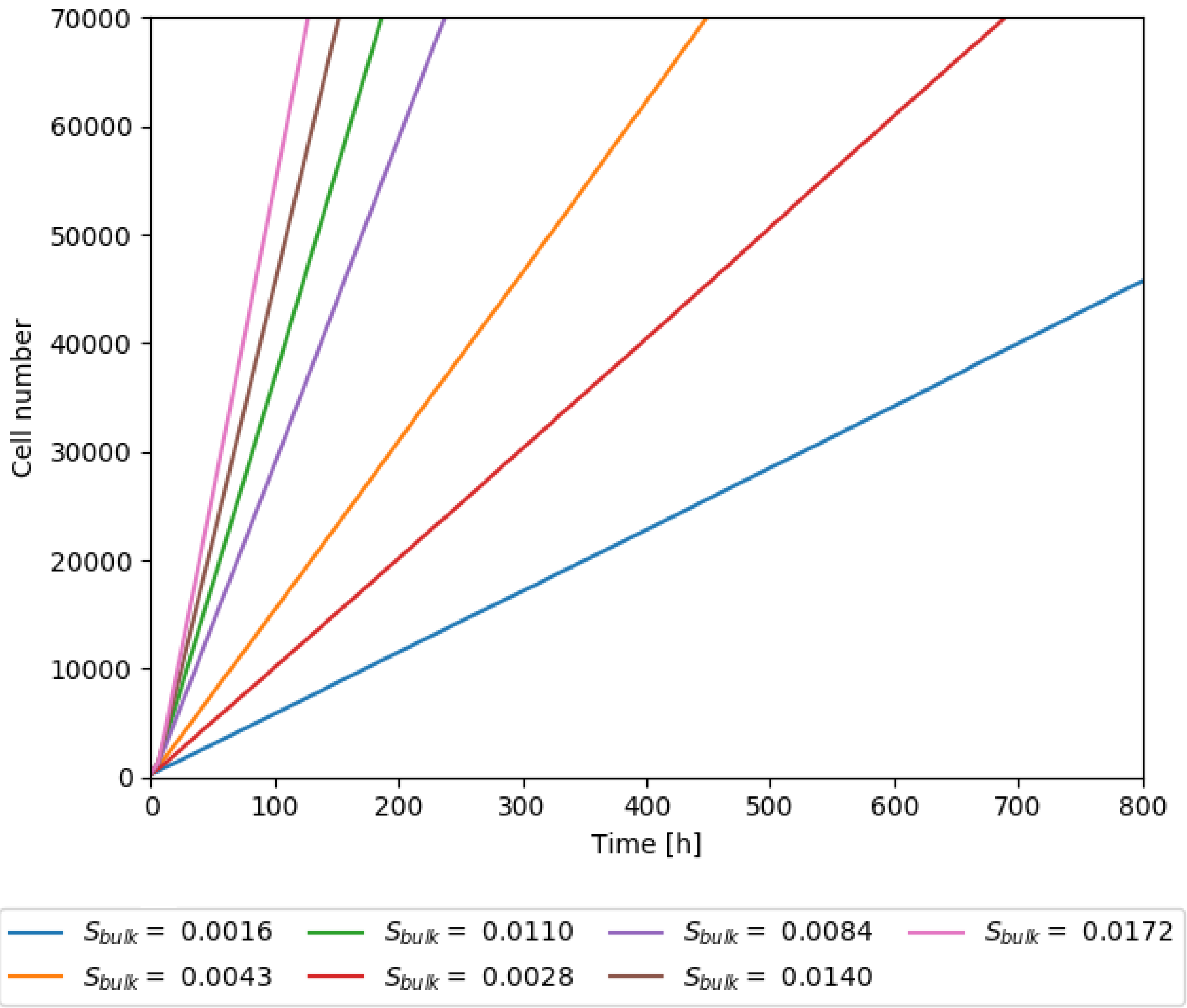


**Figure S1. Relationship between time and cell number.** This Figure shows the linear relationship between time and cell number for simulations with varying bulk nutrient concentration $S_{bulk}$. Input parameters are otherwise as specified in Table I of the main text. As detailed in the text, in the rest of our plots we chose to use cell number as a proxy for time, with the overall growth rate being the conversation factor between the two.

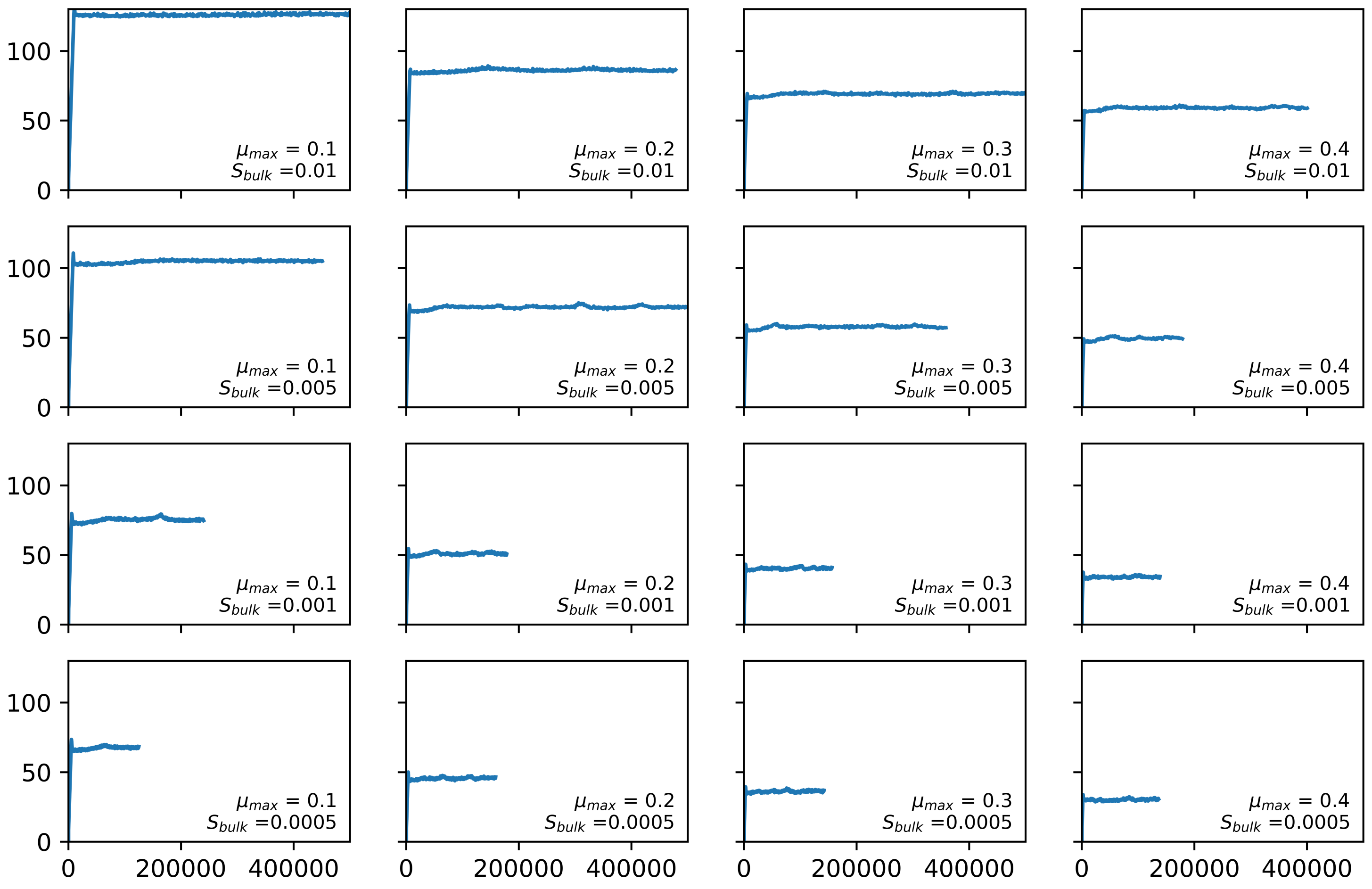


**Figure S2. Mean active layer thickness trajectories.** This figure shows the trajectories of the mean active layer thickness from our grid of simulations, varying $\mu_{max}$ and $S_{bulk}$. Units of $\mu_{max}$ are $h^{-1}$ and units of $S_{bulk}$ are g/L.

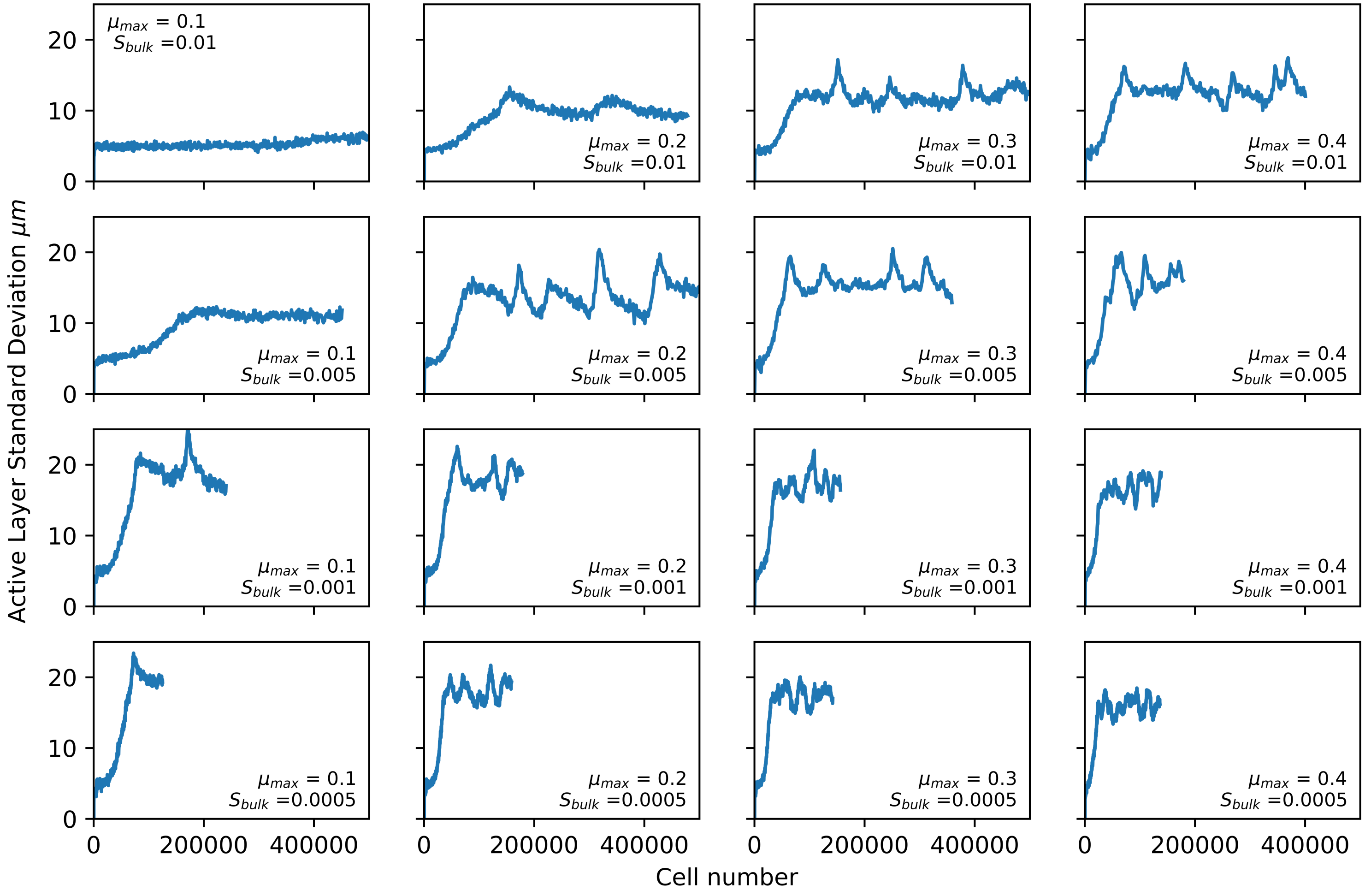


**Figure S3.** **Standard deviation of the active layer thickness trajectories.** This figure shows the trajectories of the standard deviation of the active layer thickness for our grid of simulations, varying $\mu_{max}$ and $S_{bulk}$. Units of $\mu_{max}$ are $h^{-1}$ and units of $S_{bulk}$ are g/L.

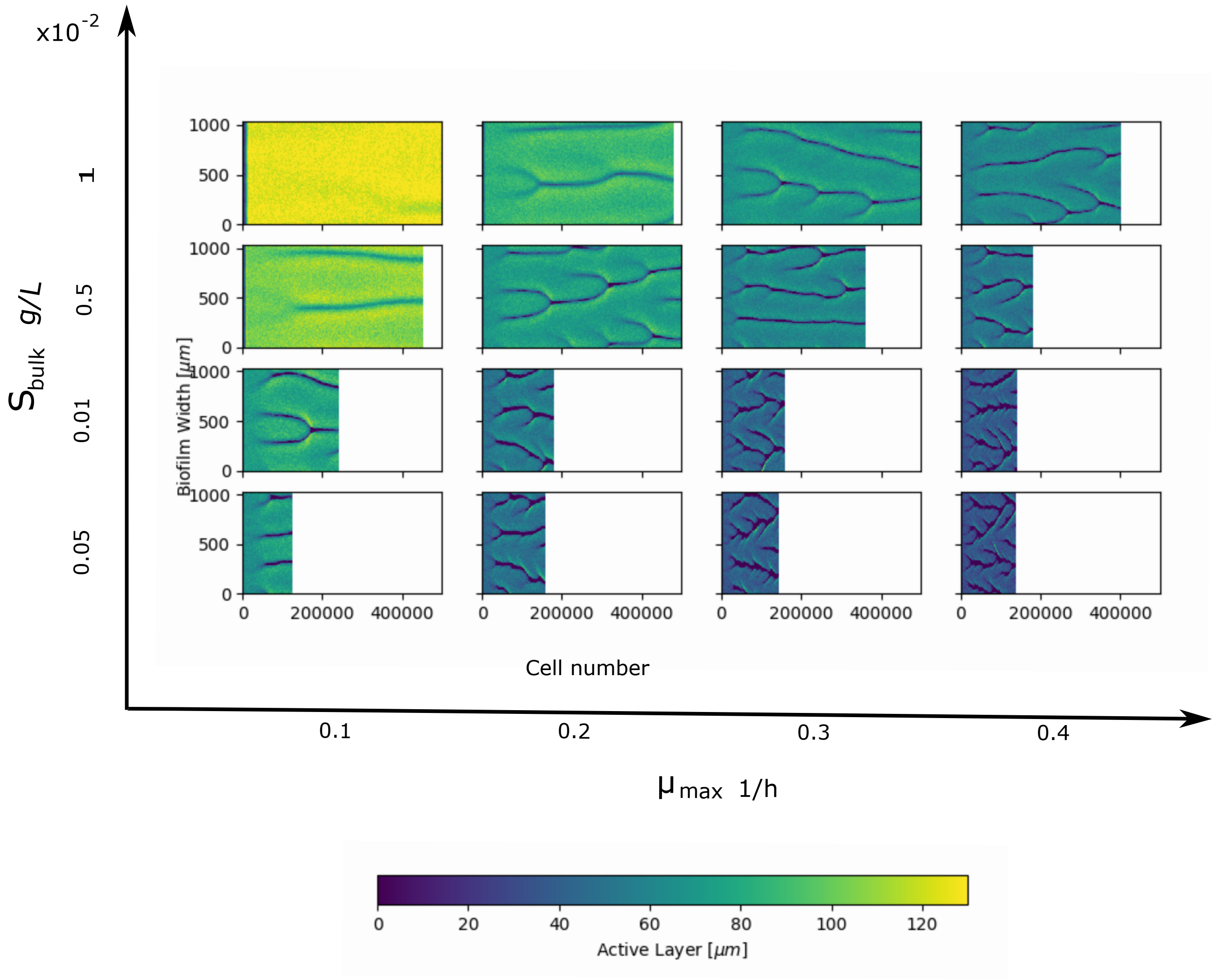


**Figure S4. Active layer gap dynamics.** This Figure shows 'kymographs' of the local active layer thickness for our grid of simulations with varying maximum specific growth rate $\mu_{max}$ and maximum nutrient concentration $S_{bulk}$, as outlined in Figure 2 of the main text. The other input parameters for the simulations are as in Table I of the main text. For each subplot, the x-axis is cell number, the y-axis is the position along the biofilm width and the colour shows the local active layer thickness. As we saw in Figure 5 of the main text, the dark lines on the kymographs correspond to the motion of active layer gaps. However, it is important to note that our two dimensional biofilms have been collapsed into one dimension in these plots. This means that in the kymographs for the pinned phase (for example, $\mu_{max} = 0.3$ 1/h, $S_{bulk} = 0.001$ g/L), the apparent merging of many small active layer gaps with the larger active layer gap are in fact 'secondary' pinning sites which appear in the original biofilm fingers - or in other words, branching behaviour (see the bottom right panels of Figure 2 in the main text).

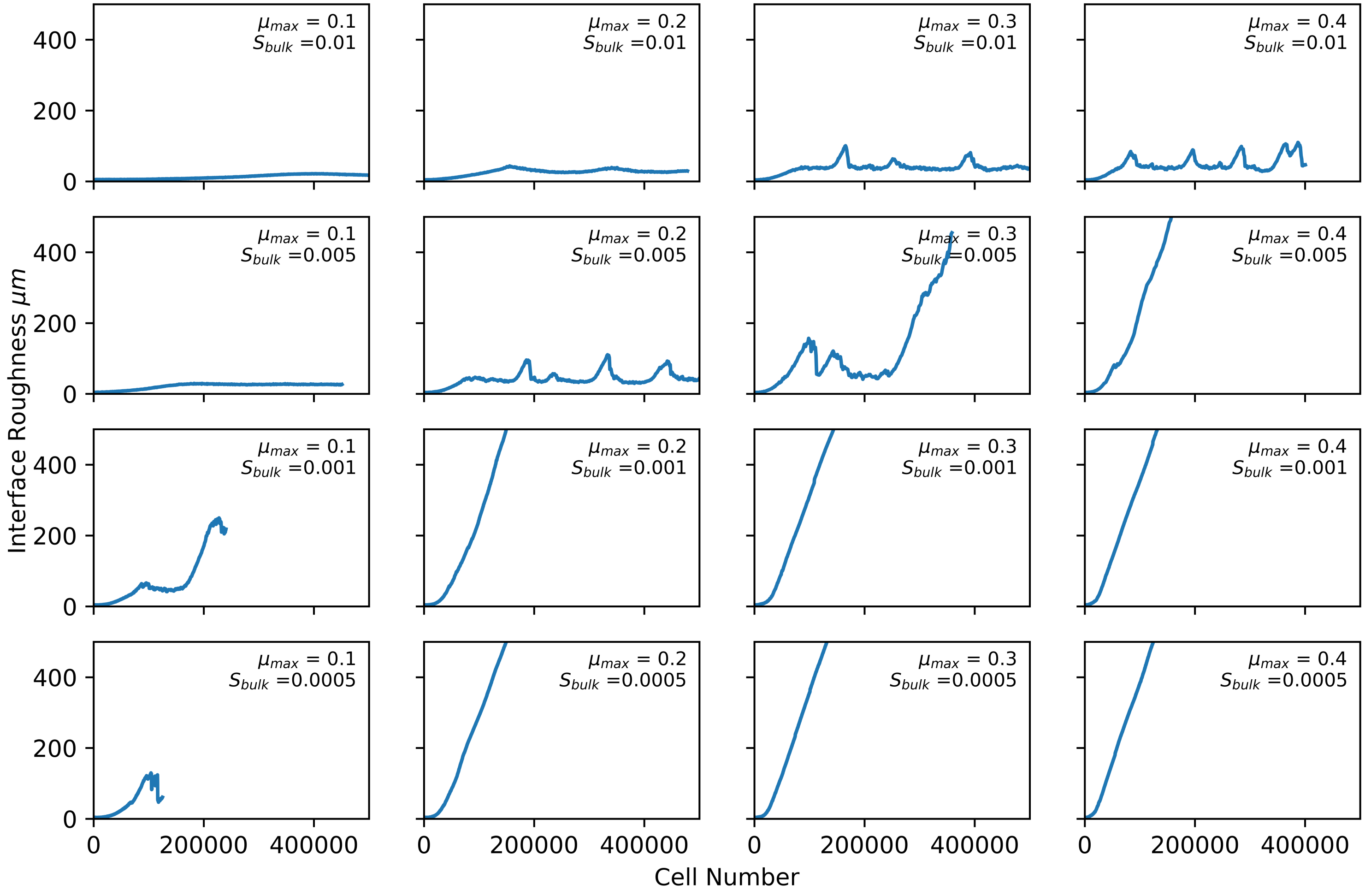


**Figure S5. Interface roughness trajectories.** This figure shows the trajectories of the interface roughness for our grid of simulations, varying $\mu_{max}$ and $S_{bulk}$. Units of $\mu_{max}$ are $h^{-1}$ and units of $S_{bulk}$ are g/L.

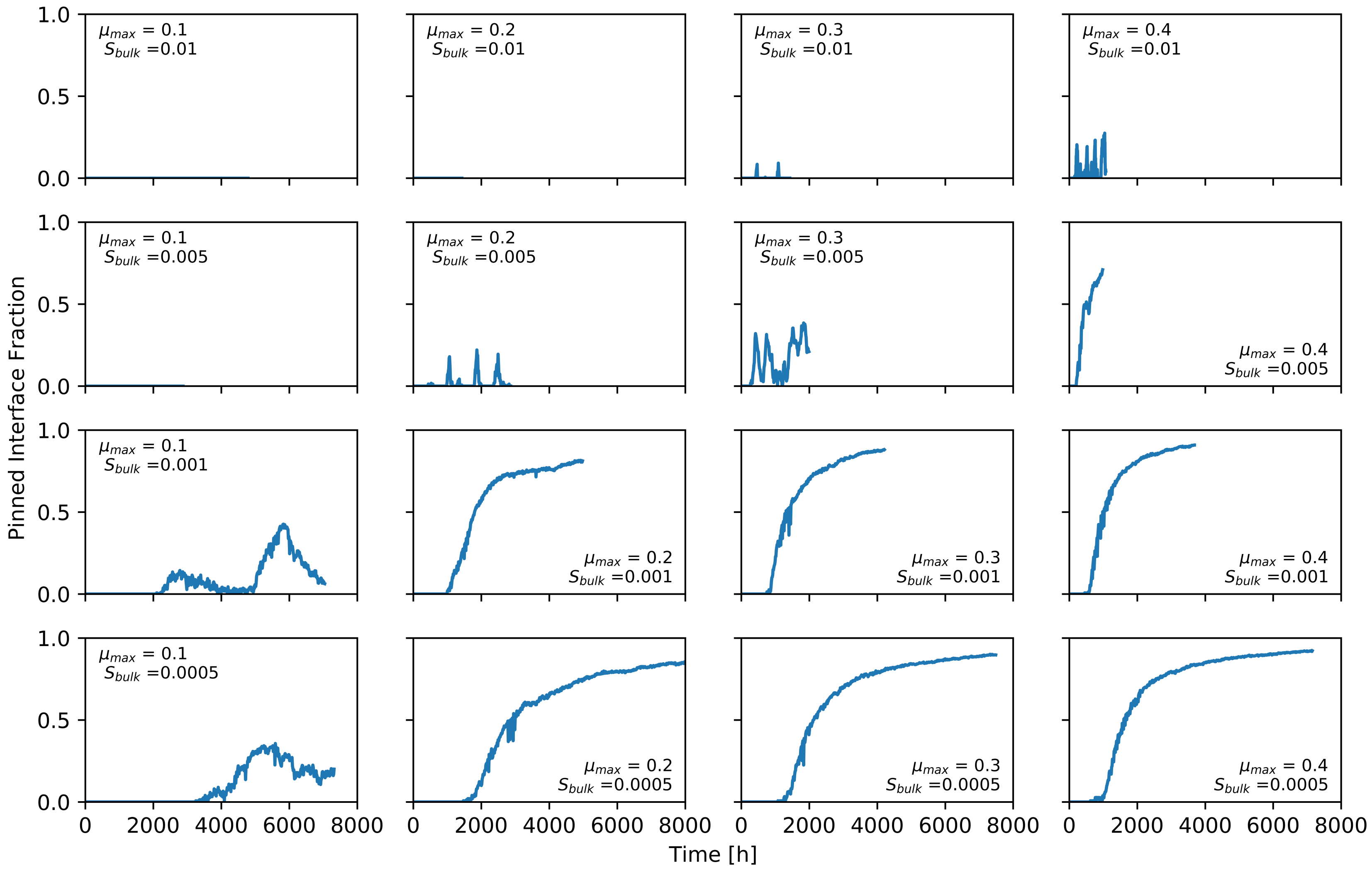


**Figure S6. Pinned interface fraction trajectories.** This figure shows the trajectories of the pinned interface fraction for our grid of simulations, varying $\mu_{max}$ and $S_{bulk}$. Units of $\mu_{max}$ are $h^{-1}$ and units of $S_{bulk}$ are g/L. Note the trajectories are plotted against time rather than cell number in this figure. As discussed in the text, computational challenges mean the steady state is not reached in all of the pinned phase simulations.

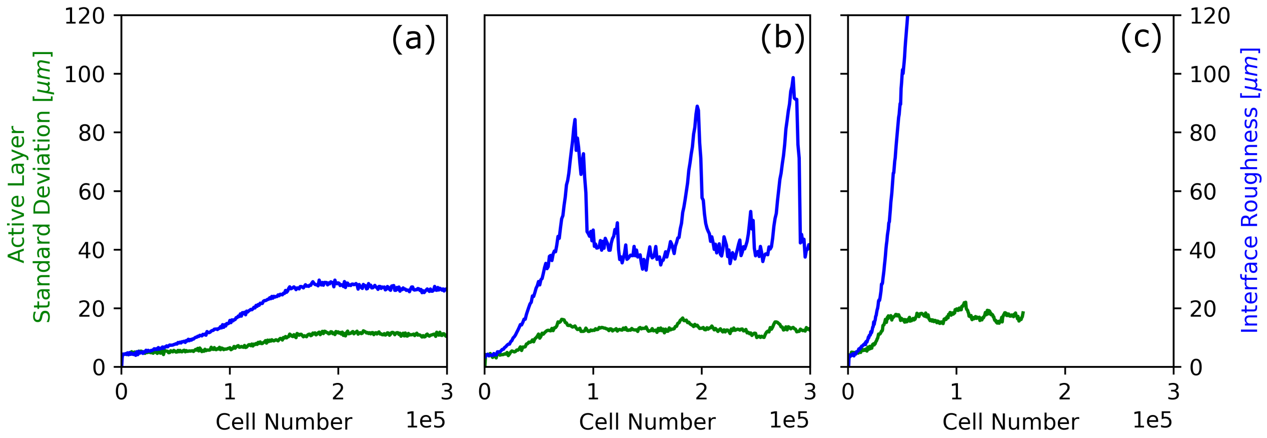


**Figure S7. Contrast between dynamics of the standard deviation of the active layer thickness and the interface roughness.** The standard deviation of the active layer thickness (green lines, in $\mu$m) is plotted on the same scale as the interface roughness (blue lines, also in $\mu$m). Panel (a) shows an example simulation in the unpinned phase ($S_{bulk} = 0.01$ g/L, $\mu_{max} = 0.1$ 1/h), panel (b) shows a simulation in the transiently pinned phase ($S_{bulk} = 0.01$ g/L, $\mu_{max} = 0.4$ 1/h) and panel (c) shows a simulation in the pinned phase ($S_{bulk} = 0.0005$ g/L, $\mu_{max} = 0.4$ 1/h). The standard deviation of the active layer thickness reaches a steady state even in simulations where the interface roughness is strongly fluctuating or monotonically increasing. Comparing with Figure 6 of the main text, panels (b)-(d), we can see that when active layer gaps, and hence pinning sites, emerge (i.e. the pinned interface fraction is non-zero), fluctuations in the active layer (standard deviation of the active layer thickness) become uncoupled from the fluctuations in the interface height (the interface roughness). In the unpinned phase (panel (a)), there is no pinning and the fluctuations in the active layer thickness and in the interface follow broadly the same trajectory. In the transiently pinned phase (panel (b)), the fluctuations follow a similar trajectory until gaps in the active layer appear (by comparison with Figure 6(c) of the main text). In the pinned phase (panel (c)), the fluctuations diverge at the point at which pinning sites arise (by comparison with Figure 6(d) of the main text).

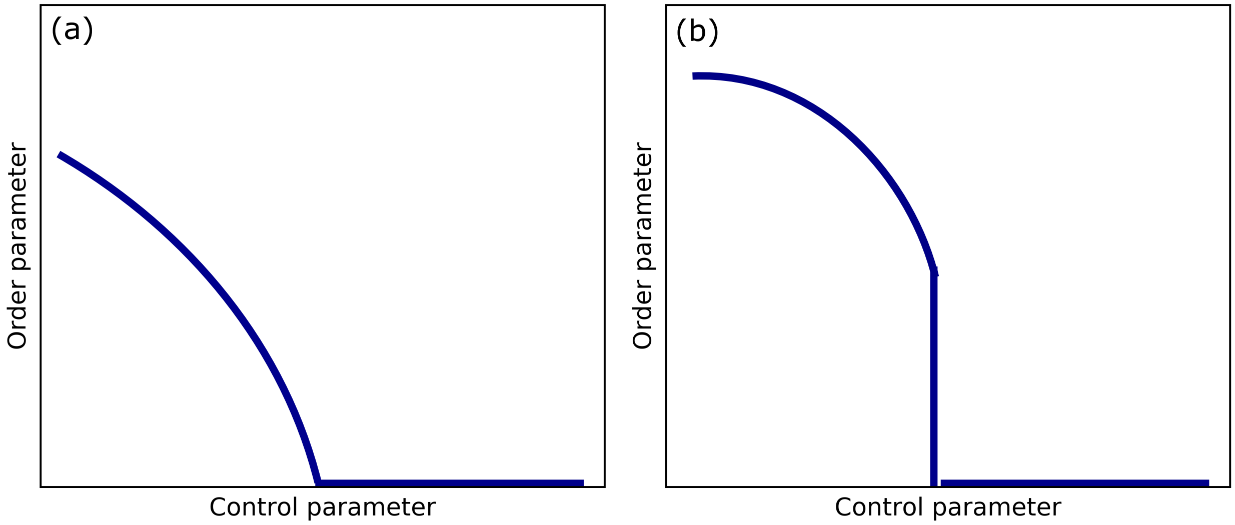


**Figure S8. Sketches of phase diagrams for continuous and discontinuous phase transitions.** In a 'continuous transition', the order parameter changes continuously from zero to a finite value as the control parameter varies (sketch (a)). In a 'discontinuous transition', the order parameter jumps discontinuously from zero to a finite value at a critical value of the control parameter (sketch (b)).

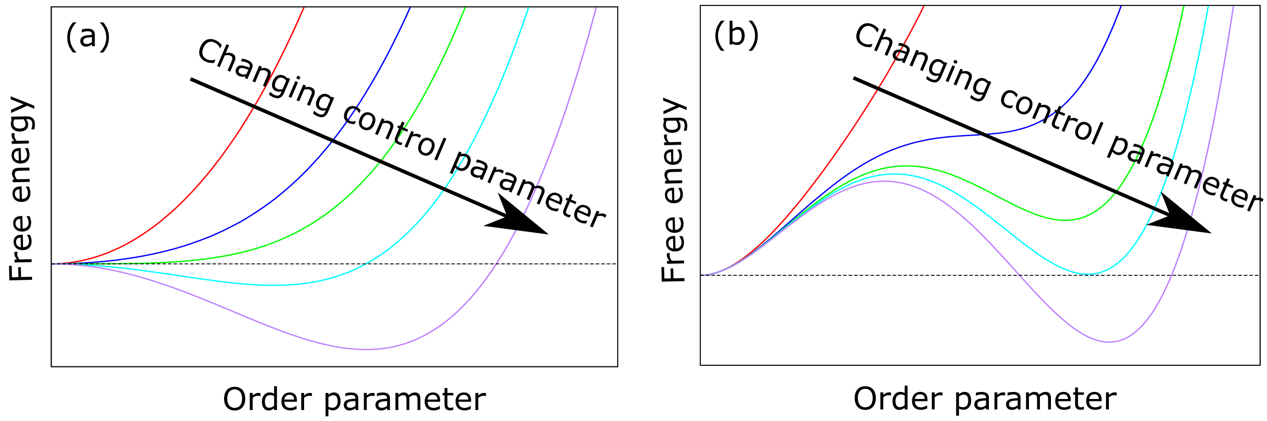


**Figure S9.** **Connecting continuous and discontinuous transitions to phase transition kinetics** The figure sketches free energy profiles for continuous and discontinuous phase transitions in an equilibrium system with a non-conserved order parameter. The coloured lines correspond to different values of the control parameter. In equilibrium systems, the minimum of the free energy profile determines the equilibrium state of the system. Sketch (a) shows a continuous phase transition. Here, the free energy profile has a single minimum. Initially (red line) the minimum is at zero on the order parameter axis, indicating that the average value of the order parameter will be zero. However as the control parameter changes, the position of the minimum shifts to the right, indicating that the average value of the order parameter becomes non-zero. The position of the minimum changes continuously as the control parameter varies. A change in control parameter will result in an immediate change in system state as the free energy minimum shifts. Sketch (b) shows a discontinuous phase transition. Initially (red line), the free energy profile has a single minimum at zero on the order parameter axis; hence the system will on average have a zero value of the order parameter. As the control parameter changes, the free energy profile develops a shoulder, and eventually a secondary minimum on the right hand side, at a non-zero value of the order parameter. This secondary minimum becomes deeper, until at a critical value of the control parameter the secondary minimum becomes the global minimum of the free energy profile. The equilibrium state of the system then shifts discontinuously from a zero value of the order parameter to the non-zero value corresponding to the secondary minimum in the free energy profile. For the discontinuous transition, because the free energy profile can have more than one minimum, there is the potential for metastability: when the control parameter changes, the system can be kinetically ‘stuck’ in a free energy minimum that is not the global minimum. To transition to the global minimum the system requires a fluctuation to overcome a free energy barrier; this phenomenon is known as nucleation. Although concepts such as free energy and metastability are ill-defined for our growing biofilms (being far from equilibrium), nucleation-like phenomena have also been observed in non-equilibrium systems.

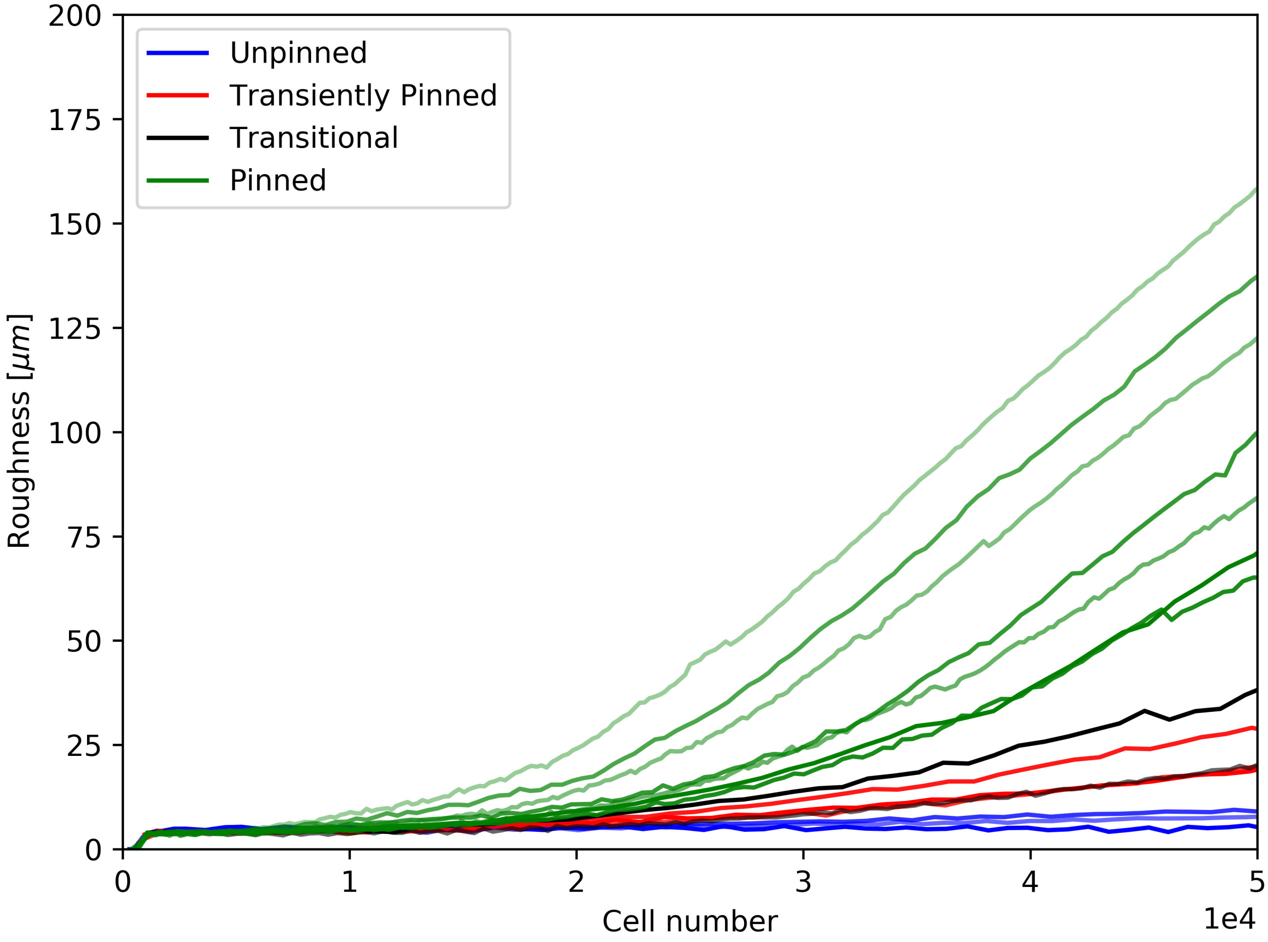


**Figure S10.** **Interface roughness at early times.** This Figure shows the early-time dynamics of the interface roughness trajectories of Figure 6(a) of the main text. In most of the simulations, there is a pseudo-steady state during the early growth behaviour (i.e. for small cell numbers $< 1 \times 10^4$). The behaviour at these early times is very different to the true steady-state behaviour that we observe at later times.

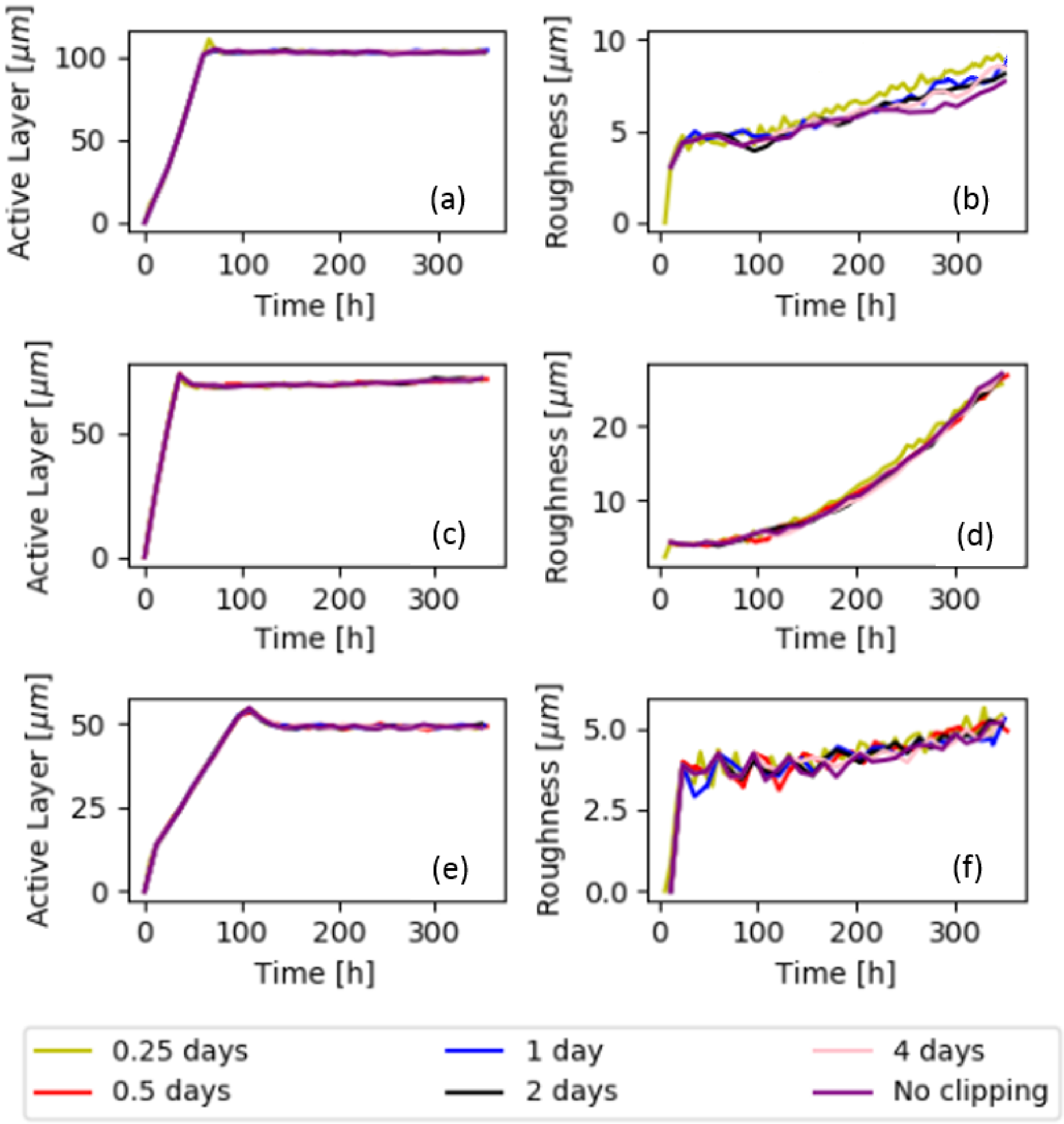


**Figure S11. Early time test case for clipping algorithm.** This figure compares the results of clipped simulations with a simulation without clipping, which is possible for short time scales. We plot trajectories of the average active layer thickness and interface roughness for test simulations with different frequencies of clipping and without clipping. Each pair of plots (a) and (b); (c) and (d); (e) and (f) are for sets of simulations with different parameter sets. The different coloured lines on these plots are for simulations with a different simulation segment length $T_s$ (i.e. a different clipping frequency; indicated in the legend) and for a continuous simulation in a single segment (i.e. no clipping). The trajectories of the active layer thickness and the interface roughness do not depend on the clipping frequency. We note that iDynoMiCS simulations are inherently stochastic, so trajectories are not expected to be exactly identical.

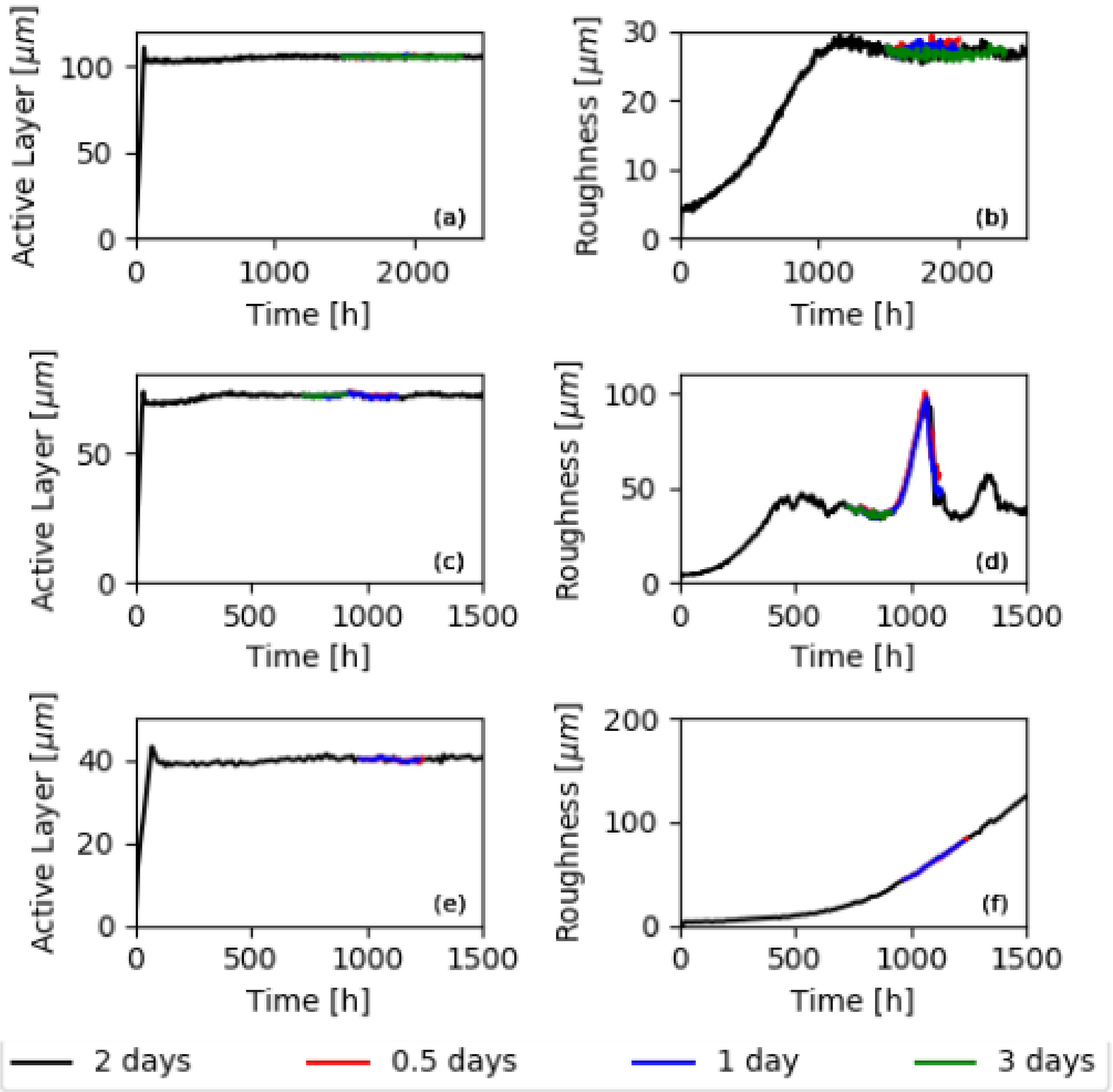


**Figure S12.** **Late time test case for clipping algorithm.** Here we compare results for long time simulations with different clipping frequencies. We plot trajectories of the average active layer thickness and interface roughness for test simulations with different frequencies of clipping. Each pair of plots (a) and (b); (c) and (d); (e) and (f) are for sets of simulations with different parameter sets. The different coloured lines on these plots are for simulations with a different simulation segment length $T_s$ (i.e. a different clipping frequency; see legend). The black line shows results from a long simulation with clipping frequency once per 2 days; new simulations with different clipping frequencies were initiated from a mid-time configuration of the black simulation. The results show that the simulation trajectories do not depend on the clipping frequency; hence we conclude that the clipping procedure does not significantly perturb the simulations.
